# Distinct cellular immune signatures in acute Zika virus infection are associated with high or low persisting neutralizing antibody titers

**DOI:** 10.1101/2021.05.27.446054

**Authors:** Elizabeth E. McCarthy, Pamela M. Odorizzi, Emma Lutz, Carolyn P. Smullin, Iliana Tenvooren, Mars Stone, Graham Simmons, Peter W. Hunt, Margaret E. Feeney, Philip J. Norris, Michael P. Busch, Matthew H. Spitzer, Rachel L. Rutishauser

**Affiliations:** Departments of Otolaryngology-Head and Neck Surgery and of Microbiology and Immunology, University of California San Francisco; San Francisco, CA, USA; Department of Medicine, Zuckerberg San Francisco General Hospital, University of California San Francisco; San Francisco, CA, USA; Vitalant Research Institute; San Francisco, CA, USA; Department of Laboratory Medicine, University of California San Francisco; San Francisco, CA, USA; Department of Pediatrics, University of California San Francisco; San Francisco, CA, USA; Parker Institute for Cancer Immunotherapy; San Francisco, CA, USA; Chan Zuckerberg Biohub; San Francisco, CA, USA

## Abstract

Although the formation of a durable neutralizing antibody response after an acute viral infection is a key component of protective immunity, little is known about why some individuals generate high versus low neutralizing antibody titers to infection or vaccination. Infection with Zika virus (ZIKV) during pregnancy can cause devastating fetal outcomes, and efforts to understand natural immunity to this infection are essential for optimizing vaccine design. In this study, we leveraged the high-dimensional single-cell profiling capacity of mass cytometry (CyTOF) to deeply characterize the cellular immune response to acute and convalescent ZIKV infection in a cohort of blood donors in Puerto Rico incidentally found to be viremic during the 2015-2016 epidemic in the Americas. During acute ZIKV infection, we identified widely coordinated responses across innate and adaptive immune cell lineages. High frequencies of multiple activated innate immune subsets, as well as activated follicular helper CD4+ T cells and proliferating CD27-IgD-B cells, during acute infection were associated with high titers of ZIKV neutralizing antibodies at 6 months post-infection. On the other hand, low titers of ZIKV neutralizing antibodies were associated with immune features that suggested a cytotoxic-skewed immune “set-point.” Our study offers insight into the cellular coordination of immune responses and identifies candidate cellular biomarkers that may offer predictive value in vaccine efficacy trials for ZIKV and other acute viral infections aimed at inducing high titers of neutralizing antibodies.

**One Sentence Summary:** Mass cytometry reveals acute ZIKV infection cellular immune signatures that predict high or low neutralizing antibody titers 6 months post-infection.

## INTRODUCTION

Infection of pregnant individuals with Zika virus (ZIKV), a flavivirus primarily transmitted to humans via the bite of an infected mosquito, can lead to persistent viral replication in the placenta and fetal brain that is associated with devastating fetal neurologic outcomes (*1–4*). In contrast, for the majority of non-pregnant immunocompetent adults, ZIKV virus is rapidly cleared from the plasma (*5–9*), and infection is accompanied by mild, self-limited symptoms such as fever, rash, and joint pain or can be asymptomatic (*10, 11*). Since the recent 2015-2016 epidemic in the Americas, there has been a considerable effort towards the development of a ZIKV vaccine, particularly for the prevention of mother-to-child transmission of infection (*12–16*). The majority of vaccine candidates in the pipeline aim to induce durable, high-titer neutralizing antibody responses, which have been shown to confer protection in animal models (*17, 18*).

After natural infection with ZIKV in humans, robust ZIKV-specific antibody responses are generated (*11, 12*); however, there is wide inter-individual variation in the levels of ZIKV-specific antibodies that persist in the serum (*11, 19*). Immunity to subsequent infection with ZIKV is likely to be influenced by the magnitude and durability of the ZIKV neutralizing antibody response (*12, 13, 20*), but little is known about the factors that contribute to inter-individual variability in this response. There is substantial cross-reactivity between virus-specific antibodies (*19, 21, 22*) and T cell responses (*23–25*) generated after infection with ZIKV and the closely-related and often co-circulating dengue virus (DENV). However, prior DENV exposure alone does not appear to explain the wide range of ZIKV antibody titers observed after natural infection (*19*). For other pathogens, baseline immune characteristics and/or signatures of early immune responses acutely after infection or vaccination have been shown to correlate with the magnitude of pathogen-specific antibody titers (*26–33*). Some aspects of the innate cytokine and cellular immune responses to ZIKV infection have been described in humans (*34–40*). However, the relationship between the acute-phase immune response and the generation of ZIKV-specific antibodies has not been characterized. This is in part due to the inherent challenges in identifying and establishing longitudinal cohorts of individuals identified during the earliest days of the acute phase of a natural infection.

Here, we leverage the high-dimensional single-cell profiling capacity of mass cytometry (CyTOF) to deeply characterize the cellular innate and adaptive immune response during acute and convalescent ZIKV infection. We evaluated longitudinal peripheral blood samples collected from 25 individuals in a natural history cohort (REDS-III) of healthy, non-pregnant adults from Puerto Rico who were found to be viremic with ZIKV at the time of blood donation during the recent ZIKV epidemic of 2015-2016 (*9, 41, 42*). We found broadly coordinated cellular responses across immune cell lineages during acute ZIKV infection and identified distinct cellular immune signatures during acute ZIKV infection that were associated with the development and persistence of low versus high neutralizing antibody titers. Future vaccine efficacy trials for ZIKV and other acute viral infections may benefit from the inclusion of these candidate cellular biomarkers to aid in the prediction of neutralizing antibody titers.

## RESULTS

### Identifying Immune Cell Populations that Respond to Acute ZIKV Infection

To characterize the cellular immune response to acute ZIKV infection, we designed two CyTOF panels to phenotype innate immune and B cell features (panel 1) and T cell features (panel 2; **Table S1**). We used these panels to analyze viably cryopreserved peripheral blood mononuclear cells (PBMCs) collected longitudinally during acute, early, and late convalescent phases of infection from 25 otherwise healthy blood donors in Puerto Rico who were incidentally found to be viremic for ZIKV at the time of blood donation (“index visit”; study participants are part of the larger REDS-III cohort; **Fig. 1A**, **Table S2** and **Data file S1**). In this cohort, 28% (7 of 25) of the participants were female, and the median age was 45 years (range 21-71). Longitudinal samples at all three timepoints were available from 19 of the 25 participants. At the first (“acute”) PBMC collection timepoint, 40% (10 of 25) of the donors were symptomatic with acute ZIKV infection, and all participants mounted a detectable ZIKV IgM, IgG, and neutralizing antibody response (reported as the 80% neutralization titers: NT_80_; **Fig. 1B**). Although all participants were viremic at the index visit, 68% (17 of 25) had not yet formed ZIKV-specific IgM responses, indicating that they were likely enrolled within the first 8 days of infection (*41*). Of the participants with a collection visit at the first (“acute”) PBMC collection timepoint (median 8 [range 5-12] days after index visit), 100% (23 of 23) had formed IgM antibodies and 22% (5 of 23) had residual detectable plasma viremia. There was substantial variation in both peak neutralizing antibody titers (ZIKV NT_80_ titers: 84 - 37,872) and follow-up titers 6 months after the index visit (0 - 6,286).

**Fig. 1.**
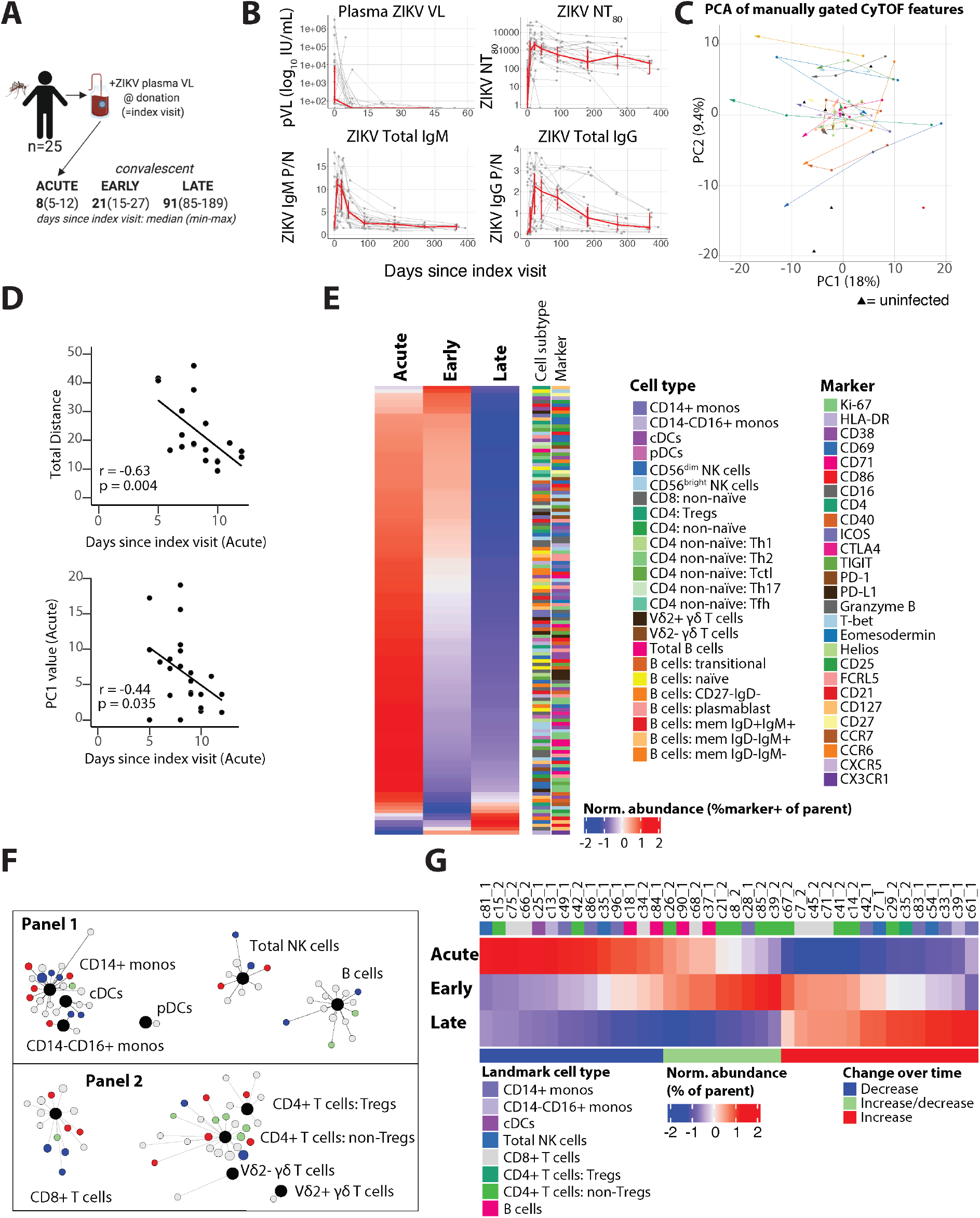
Acute infection with ZIKV elicits profound phenotypic changes across peripheral blood cellular immune populations. (A) 25 adults viremic with acute ZIKV infection at the time of blood donation (“index visit”) had peripheral blood sampling at up to three timepoints: during the acute phase of infection, early and/or late convalescence. (B) Plasma ZIKV viral load (VL), neutralizing antibody titers (NT_80_), and total IgG and IgM levels of study cohort participants. Red line connects median values at each sampling timepoint (+/− 95% Confidence Interval, CI). (C) Line plots for each participant in PCA space with arrows indicating direction from early to later timepoints. Black triangles denote coordinates for 8 uninfected control samples. (D) Scatterplots of days since index visit at the acute timepoint and the value of PC1 at the acute timepoint or the total distance traveled in PCA space between the acute and late convalescent timepoints (Spearman’s correlation with regression line). (E) Heatmap showing the z-score normalized frequency of the log-adjusted feature abundances for the manually gated phenotypic features that change significantly from the acute to the late convalescent timepoint. (F) SCAFFoLD maps showing clusters of cells associated with landmark cell population nodes (black dots). Clusters that significantly change in abundance between the acute and late convalescent timepoints based on LME model fit are denoted in the following colors: increase (red), decrease (blue), or increase and then decrease (green). (G) Heatmap showing the normalized abundance of the clusters (z-score based on % of parent cell type population) that change significantly. Significance in (E-G) based on linear mixed effects (LME) model fit with Benjamini-Hochberg false discovery rate (FDR)-corrected p-value, p_adj< 0.05.

We first sought to characterize how acute ZIKV infection perturbs the frequency and phenotype of different immune cell types in peripheral blood. We manually gated major landmark immune cell populations defined by standard lineage markers (e.g., classical [CD14+] monocytes, non-classical [CD14-CD16+] monocytes, plasmacytoid dendritic cells [pDCs], classical DCs [cDCs], CD56^bright^ and CD56^dim^ NK cells, CD4+ T cells, CD8+ T cells, B cells, etc.) and classically-defined adaptive immune cell subsets (see **Fig. S1** for gating strategy and **Table S1** for mass cytometry antibodies used). We first evaluated the relative abundance of 40 cell types consisting of landmark immune cell populations (as a proportion of total live CD45+ cells) and adaptive immune subsets (as a proportion of their parent landmark population). We then evaluated the Boolean expression of 30 different phenotypic surface and intracellular proteins on these parent cell types, which yielded a total of 286 unique phenotypic features (see **Table S3** for phenotypic feature gating strategy and **Data file S2** for raw data of all cell type frequency and phenotypic features).

To broadly determine how the immune state is perturbed in the context of ZIKV infection, we performed principal component analysis (PCA) on the manually-gated cell type frequency and phenotypic features identified from our CyTOF analysis. Features were adjusted for age and sex (median +/− standard deviation contribution to the variance for individual features: 1.63 +/− 5.83%, 1.17 +/− 3.16%, respectively; age/sex contribution to variance for individual features can be found in **Data file S2**). We mapped the trajectories across the three timepoints in PCA space, as defined by the first two principal components (PC1 and PC2) for each individual ZIKV-infected participant (**Fig. 1C**) as well as 8 control ZIKV-uninfected blood donors sampled from the same blood donation sites prior to the ZIKV epidemic (black triangles). While there was variation between individuals, most participants followed a similar general trajectory from right-to-left along PC1 as they progressed from acute to convalescent ZIKV infection. The number of days between the index and the acute timepoint negatively correlated with the total distance travelled in PCA space across the top five PCs as well as the value of PC1 at the acute timepoint (Spearman’s r=-0.63, p=0.004; r=-0.44, p=0.035, respectively; **Fig. 1D**). These correlations suggest that both parameters (the PC1 coordinate and the distance travelled) correspond to movement in “virtual” infection space as participants resolve their ZIKV infection. Of note, we did not observe coordinated trajectories in PCA space across longitudinal sampling of 6 ZIKV-uninfected control individuals (**Fig. S2**).

In order to understand which cellular features contribute to this coordinated movement over time, we used linear mixed effect modelling on the above age- and sex-adjusted feature abundances (see **Data file S3** for tables of all statistical analyses performed). We first determined whether the abundance of major landmark populations and adaptive immune cell subsets changed significantly over time since index visit. We were surprised to find that the frequency of most of these immune cell types did not change significantly across the three sampled timepoints, with the exception of small changes in the abundance of total B cells and several CD4+ T cell subsets (naïve, non-naïve, TEM, Treg, Th1, Th17, Tfh/Th17; **Fig. S3**).

We next analyzed the 286 phenotypic features identified by manual gating and found that 128 changed significantly across the three sampled timepoints (p_adj<0.05; **Fig. 1E**). Of note, when we analyzed cross-sectional data from the acute timepoint versus the 8 uninfected participants, only 53 features were significantly different (p_adj<0.05), reflecting the power of our longitudinal paired analysis to identify immune features that are altered by acute infection even with the inherent inter-individual variation in human datasets. The vast majority of these changing features (122 of 128, 95%) decreased in abundance between the acute and late convalescent timepoints. A subset of these features remained elevated or even increased at the early convalescent timepoint (e.g., several TIGIT+ CD4+ T cell populations, and several CD69+ B cell populations), while others decreased sharply between the acute and early convalescence stage (e.g., most populations expressing Ki-67 and CD71). Of note, Helios+ Tregs, CD127+ non-naïve CD8+ T cells and NK cells, multiple CD21+ memory B cell subsets, and CX3CR1+ non-classical monocytes were the only features whose frequencies (as a proportion of the parent population) significantly increased over time. We observed no significant changes over time in the same set of manually gated features in longitudinal analysis of 6 uninfected controls (**Data files S2** and **S3**).

To leverage the richness of our high dimensional single cell dataset, we performed unsupervised clustering of our dataset using the SCAFFoLD algorithm that we have described previously (*43, 44*). This algorithm performs CLARA clustering with the phenotypic markers from the mass cytometry panel to partition the cells into a user-defined number of clusters and then creates a force-directed graph to connect those clusters to user-defined landmark reference cell populations. We observed high concordance in the frequency of the pre-defined landmark immune cell populations between our manual gating and unsupervised clustering approaches (**Fig. S4**). We applied the same linear mixed effect model approach to the cluster abundances from the SCAFFoLD algorithm and found that 15 of 34 clusters assigned to innate immune cell types and 23 of 56 clusters assigned to adaptive immune cell types (innate immune and B cell clusters from CyTOF Panel 1, T cell clusters from Panel 2) changed significantly in abundance over time between the acute and late convalescent phase of ZIKV infection (**Fig. 1F, G**; full raw and statistical analysis data available in **Data files S2** and **S3**). Similar to our prior analysis, we observed diversity in the direction and speed with which clusters changed in abundance over the three timepoints.

### Innate Immune Cell Activation in Acute ZIKV Infection

Little is known about the innate immune response to acute ZIKV infection in humans. CD14+CD16+ monocytes have been shown to increase in the peripheral blood of children with acute infection and are themselves a major target for ZIKV infection (*37, 45*). We also observed an increase in intermediate (CD14+CD16+) monocytes during acute ZIKV infection followed by a decline in early convalescent phase (median 1.0% to 0.27% of live cells, p_adj=0.002; **Fig. 2A, B**). The proportion of intermediate monocytes expressing Ki-67, CD69, CD71, CD4, and PD-1 decreased over time, suggesting that this population was more activated during acute infection (**Fig. 2C**). We observed a decrease in the proportion of both classical (CD14+) monocytes (which includes the intermediate monocytes) and non-classical (CD14-CD16+) monocytes expressing these and additional activation and differentiation markers (**Fig. 2D**).

**Fig. 2.**
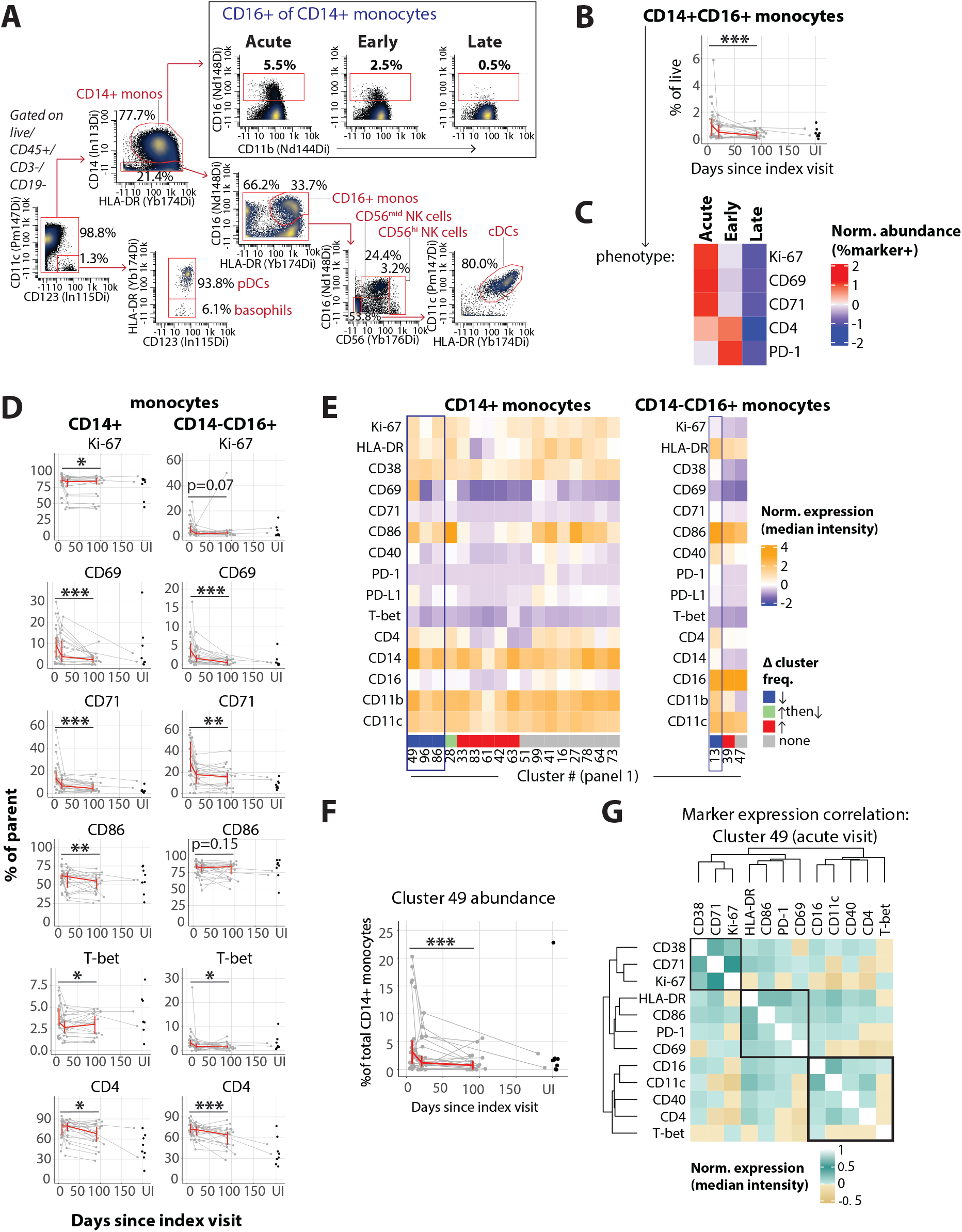
Coordinated activation marker expression on monocytes during acute ZIKV infection. (A) Gating scheme for innate immune cell types, including intermediate monocytes (CD14+ monocytes that express CD16). Percentages shown are % of parent populations in the plotted sample. (B) Frequency (as a % of total live cells) and (C) phenotype (z-scored proportion of cells that express each marker) of CD14+CD16+ monocytes across the course of acute and resolving ZIKV infection. (D) Expression of individual activation markers on CD14+ and CD14-CD16+ monocytes. (E) Heatmap showing z-score normalized median expression of indicated markers (rows) for each monocyte-associated cell cluster (column). Column annotation indicates clusters that that significantly decrease (blue), increase (red), increase and then decrease (green), or remain unchanged (grey) in abundance (as a % of the parent monocyte population; p_adj<0.05). (F) Change in abundance of CD14+ monocyte cluster 49 (as a % of CD14+ monocytes; p_adj=0.0002). (G) Spearman’s correlation matrix of marker expression on single cells in CD14+ monocyte cluster 49 from acute visit samples. p_adj values obtained by LME model fit with Benjamini-Hochberg FDR correction. *p_adj<0.05, **p_adj<0.01, ***p_adj<0.001. B, D, F: Red line connects median values at each sampling timepoint (+/− 95% CI). UI=uninfected.

Unsupervised clustering analysis **(Fig. 2E**) demonstrated that many of these activation markers (e.g., Ki-67, CD69, CD86, and CD4) were highly expressed on some of the clusters that decreased in frequency over time (identified by blue box and bar under the column). Other activation markers (e.g., PD-1, CD71) had similar expression between the decreasing and increasing clusters (**Fig. 2E**). We investigated the co-expression of these activation markers and other phenotypic markers on individual cells contained within cluster 49, a CD14+ monocyte cluster with the greatest median relative change in frequency (−75%) from the acute to late convalescent timepoints (**Fig. 2F**). This revealed three modules of markers with coordinated expression patterns: (1) a proliferative module (Ki-67, CD71, and CD38), (2) an early activation module (HLA-DR, CD86, PD-1, and CD69), and (3) a monocyte maturation/differentiation module (CD16, CD11c, CD40, and CD4; **Fig. 2G**). Interestingly, the expression of most of the markers in the maturation/differentiation module were inversely correlated with the expression of Ki-67. Thus, unsupervised analysis allowed us to identify clusters of CD14+ monocytes that appear to occupy distinct differentiation states.

The expression of activation markers (CD69, CD71, CD86, CD40, Ki-67, and HLA-DR) also significantly decreased on the cDCs over time (**Fig. S5A**). In the unsupervised clustering analysis, the cluster of cDCs with the greatest relative decrease in abundance had the highest expression of several of those markers (CD69, CD86, CD71, and CD40) compared to the clusters of cDCs whose frequencies were unchanged, suggesting that these markers are co-regulated on a population of DCs during response to acute ZIKV infection (**Fig. S5B**). The expression of several activation markers (CD38, CD86, and Ki-67) also decreased on pDCs as ZIKV infection resolved (**Fig. S5A, B**).

Amongst NK cells, there was a decrease in proliferation (Ki-67) and activation (CD38, CD69, CD71) in both CD56^bright^ and the more cytotoxic CD56^dim^ NK cells (**Figs. S5C, D**). Additionally, CD56^dim^ NK cells uniquely exhibited a decrease in the cells expressing both Granzyme B and TIGIT, with a concurrent increase in the proportion of cells expressing CD127. In contrast, CD56^bright^ NK cells also demonstrated a decrease in cells expressing CD86 and CD16 as ZIKV infection resolved. Collectively, these data demonstrate that acute ZIKV infection is characterized by the accumulation and activation of innate immune cells.

### T Cell Response to Acute ZIKV Infection

Although we did not observe a higher frequency of total or even non-naïve CD8+ T cells in acute compared to convalescent ZIKV infection, we did see a profound accumulation of non-naïve CD8+ T cells co-expressing the activation markers HLA-DR and CD38 during acute infection (median 11.3% of the non-naïve population, range 2.3-47.3%, compared to a median of 1.7% [0.4-4.5%] during late convalescent infection; **Fig. 3A, B**). The population of HLA-DR+CD38+ CD8+ T cells is thought to be enriched for antigen-specific CD8+ T cells in acute infections (*46–48*). Indeed, a median of 93.4% (range 42.4-99.7%) of the HLA-DR+CD38+ non-naïve CD8+ T cells in acute ZIKV infection had evidence of active proliferation as measured by expression of Ki-67 (as compared to a median of 60.5% [range 38.6-80.5%] in late convalescence). The proportion of HLA-DR+CD38+ CD8+ T cells expressing Ki-67 and other activation markers (ICOS, CTLA-4) decreased rapidly between acute infection and early convalescence, while the proportion of cells expressing markers associated with effector CD8+ T cell differentiation (Granzyme B and T-bet) decreased more gradually across the time course to late convalescence (**Fig. 3C**). Interestingly, there was a small but significant increase in the proportion of HLA-DR+CD38+ CD8+ T cells expressing the co-inhibitory receptor PD-1 between acute infection and early convalescence that was maintained into late convalescence (median 19.5% => 24.6% => 24.8%; p_adj=0.002). There was also a concurrent increase in cells expressing the transcription factor Helios, the expression of which is often associated with CD4+ Treg cells (**Fig. 3C**).

**Fig. 3.**
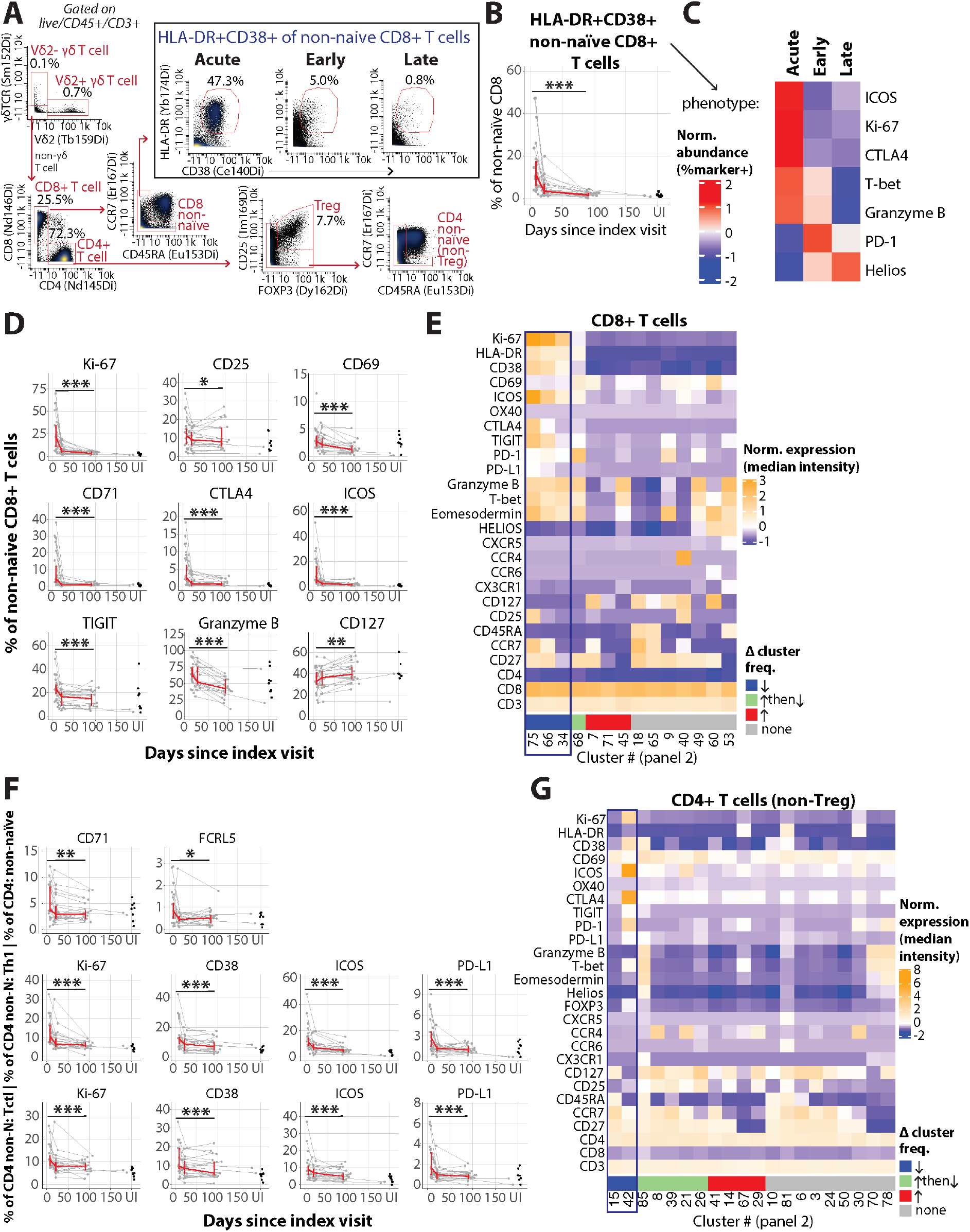
Accumulation of activated T cells during acute ZIKV infection. (A) Gating scheme for T cell subsets, including non-naïve CD8+ T cells that co-express HLA-DR and CD38 (see **Fig. S1** for full CD4+ subset gating). Percentages shown are % of parent populations in plotted sample. (B) Frequency (as a % of non-naïve CD8+ T cells) and (C) phenotype (z-scored proportion of cells that express each marker) of HLA-DR+CD38+ non-naïve CD8+ T cells across the course of acute and resolving ZIKV infection. Expression of individual activation markers on (D) non-naïve CD8+ T cells and (F) non-naïve, Th1, and Tctl subsets of CD4+ T cells. Phenotype (z-scored median expression of each marker) of (E) CD8+ T cell or (G) CD4+ T cell clusters that significantly decrease (blue), increase (red), increase and then decrease (green), or remain unchanged (grey) in abundance (as a % of the parent T cell population; p_adj<0.05). p_adj values obtained by LME model fit with Benjamini-Hochberg FDR correction. *p_adj<0.05, **p_adj<0.01, ***p_adj<0.001. B, D, F: Red line connects median values at each sampling timepoint (+/− 95% CI).

Of the total non-naïve CD8+ T cell population, there was a significant decrease following the acute phase in the proportion of proliferating (Ki-67+) cells, as well as cells expressing several activation markers (CD25, CD69, CD71, CTLA-4, PD-L1, PD-1, ICOS, TIGIT) and effector markers (Granzyme B, Eomesodermin, T-bet). Concurrently, there was an increase in the proportion of CD8+ T cells expressing CD127 (**Fig. 3D**). With our unsupervised clustering analysis, we found three clusters of non-naïve (CD45RA-) effector-differentiated (Granzyme B+T-bet+) CD8+ T cells that decreased in abundance (as a fraction of the non-naïve CD8+ T cell population) between the acute and late convalescent timepoints (summed median frequency of c75, 66, 34: 7.1% => 3.1% => 1.6%; **Fig. 3E**). There were several other dynamic changes in cluster abundance over time, including one effector-differentiated cluster (cluster 68) that increased and then decreased in abundance (median frequency: 1.4% => 1.5% => 0.63%), as well as three clusters that significantly increased in abundance between the acute and late convalescent timepoints (summed median frequency of c7, 71, 45: 21% => 30% => 33%; **Fig. 3E**). Interestingly, the cells contained within the decreasing clusters expressed multiple activation markers (e.g., Ki-67, HLA-DR, CD38, ICOS, CTLA-4, and TIGIT; cluster 68 also expressed the highest level of PD-1; **Fig. 3E**), suggesting that the expression of these markers is coordinately regulated on small sub-populations of cells during acute ZIKV infection.

Although there was not a significant change in the frequency of total CD4+ T cells between the acute and convalescent timepoints (**Fig. S2A**), we observed a decrease after the acute timepoint in the frequency of naïve, regulatory (Treg; FOXP3+CD25+), and non-naïve (non-Treg) Th1, Th17, and Tfh/Th17 CD4+ T cells and an increase in effector memory and non-naïve CD4+ T cells (**Fig. S2B**; see **Fig. 3A** and **Fig. S1** for gating). Acute ZIKV infection was associated with profound activation of the total non-naïve CD4+ T cell compartment as well as the non-naïve Th1, cytotoxic (Tctl), and Treg subsets (**Figs. 3F**, **S6A**). We did not observe a significant change in the frequency of circulating T follicular helper cells (Tfh), although some individuals in the cohort appeared to have an expansion of Tfh cells at the early convalescent timepoint (median 21 days after index visit), followed by a contraction during late convalescence (**Fig. S2B**). Similar to CD4+ and CD8+ T cells, cycling, activated, and cytotoxic non-naïve γδ T cells were present at higher frequencies during acute ZIKV infection (**Fig. S6B**).

### B Cell Response to Acute ZIKV Infection

We observed a significant increase in total B cells during the resolution of acute ZIKV infection; however, similar to most major T cell subsets, we did not observe a change in the abundance of the major B cell subsets (as a proportion of total B cells; **Fig. S2B**). We identified B cell populations enriched for cells that were actively secreting antibodies in two ways: (1) based on the expression of markers classically used to identify plasmablasts (CD38^hi^CD20^neg^, **Fig. 4A** and **Fig. S2B**), and (2) based on the expression of markers that have been used to identify “Antibody Secreting Cells” (ASCs) in Ebola infection in humans, CD71^high^CD20^neg^ non-naïve B cells (**Fig. 4A;** (*49*); there is a significant correlation in the frequency of these two populations, r=0.92 p=4E-30, **Fig. S7A**). We noted a significant decrease in the frequency of these cells between acute infection and early/late convalescence, gated as either plasmablasts or ASCs (both gated as a % of non-naïve B cells; p=0.03 and p=0.04, respectively). Phenotypically, during acute infection compared to early and late convalescence, a larger proportion of ASCs expressed the transcription factor T-bet, which has been associated with B cell responses to viral infections (*50*) (median 15.4% => 8.2% => 7.7%, p_adj=0.02; **Fig. 4C**). We also observed higher expression of T-bet in naïve and some class-switched memory B cell subsets during acute ZIKV infection (**Fig. 4D**).

**Figure 4.**
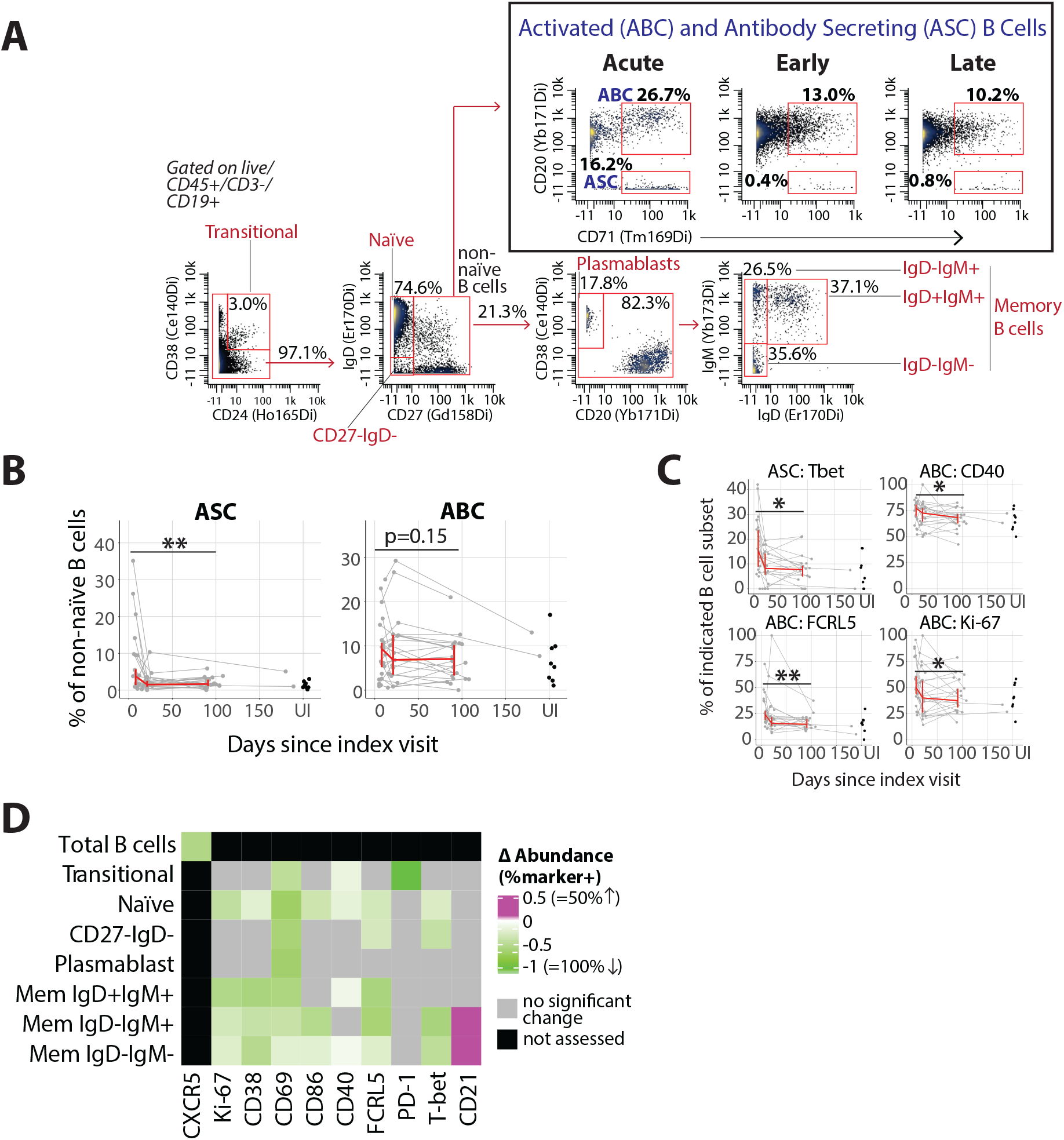
Accumulation of activated B cells during acute ZIKV infection. (A) Gating scheme for B cell subsets, including activated and antibody secreting B cells (ABC and ASC, respectively). Percentages shown are % of parent populations in the plotted sample. (B) Frequency (as a % of non-naïve B cells) and (C) phenotype of ABCs and ASCs across the course of acute and resolving ZIKV infection. (D) Relative change in frequency [(late convalescent – acute)/acute] of the expression of individual activation markers on B cell subsets (median shown). Colors indicate markers with a significant (p_adj<0.05) change in the percent of the parent population that expresses the marker are noted (increase=pink, decrease=green). p_adj values obtained by LME model fit with Benjamini-Hochberg FDR correction. *p_adj<0.05. For B, C: Red line connects median values at each sampling timepoint (+/− 95% CI).

An increased frequency of “Activated B Cells” (ABCs), as defined by high expression of both CD20 and CD71 on non-naïve B cells, has also been described in other acute infections in humans (*49, 51*). Although some individuals with acute ZIKV infection had a higher frequency of ABCs at acute or early convalescent timepoints, we did not observe this across all participants (**Fig. 4B**). We did observe a significant decrease over time in the proportion of ABCs expressing Ki-67, FCRL5, and CD40 (**Fig. 4C**). Furthermore, during acute ZIKV infection, we observed a higher proportion of B cells in multiple subsets (including naïve B cells) that were activated based on the expression of proteins other than CD71, including Ki-67, CD38, CD69, CD86, CD40, and FCRL5 (**Fig. 4D**). Based on our clustering analysis, it appears that most of these activation markers were co-regulated on two clusters of B cells whose frequency declined with the resolution of acute infection (**Fig. S7B**). A higher proportion of total B cells also expressed CXCR5 during acute infection (**Fig. 4D**). Finally, the expression of CD21 was lower on the two IgD-memory B cell populations during acute ZIKV infection (**Fig. 4D)**, which may identify cells recently exited from a germinal center (*52*).

### Coordinate Activation of Innate and Adaptive Immune Cells in Acute ZIKV Infection

We next explored how immune parameters measured during acute infection were coordinated with each other. To limit the variation in the data collected from participants during acute infection (who may have been sampled at a different number of days following infection with ZIKV), for the analysis presented in **Figures 5**-**6** we focused on 17 individuals whose anti-ZIKV IgM was negative at their index visit (which indicates that their index visit was likely within 8 days of the onset of detectable viremia (*42*)).

**Fig. 5.**
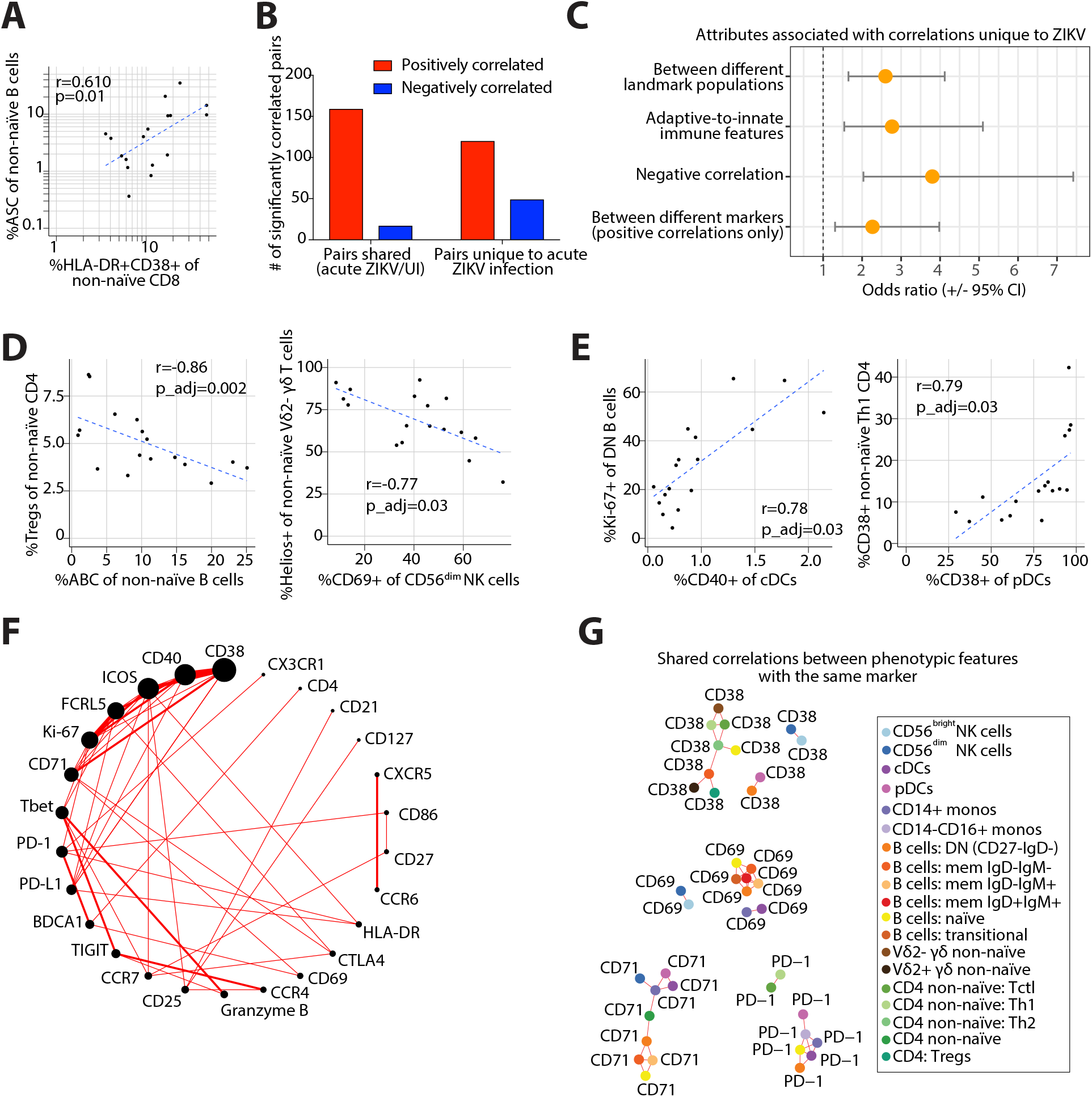
Coordinated activation across different cell types in acute ZIKV infection. (A) Scatterplot of the frequency of ASC B cells and CD38+HLA-DR+ CD8+ T cells in acute ZIKV infection with regression line. (B) Number of significant (p_adj) positive and negative correlations between cellular immune features that are present in acute ZIKV infection, grouped by those that are “unique” to ZIKV versus those “shared” with the uninfected (UI) cohort. (C) Odds ratio (+/− 95% CI) that cellular immune feature correlations unique to ZIKV infection are more likely to be associated with different correlation attributes (compared to the correlations shared with the UI cohort). Correlation plots of select features uniquely negatively (D) or positively (E) correlated in acute ZIKV infection (Spearman’s r with correltion line). (F) Circular network graph of positive correlations unique to acute ZIKV infection that are between phenotypic features with different markers. Size of node indicates number of correlations including a feature classified under the corresponding label and edge thickness is proportional to the number of correlations between features classified by the indicated nodes (markers). (G) Network graph showing of positive correlations (red edges) between phenotypic features with the same marker that are shared between acute ZIKV infection and the UI cohort. Node colors indicate cell subtype. A, D: Spearman’s correlation with regression line. B, E, F: p_adj values determined by Spearman’s correlation and adjusted with the Benjamini-Hochberg method.

**Fig. 6.**
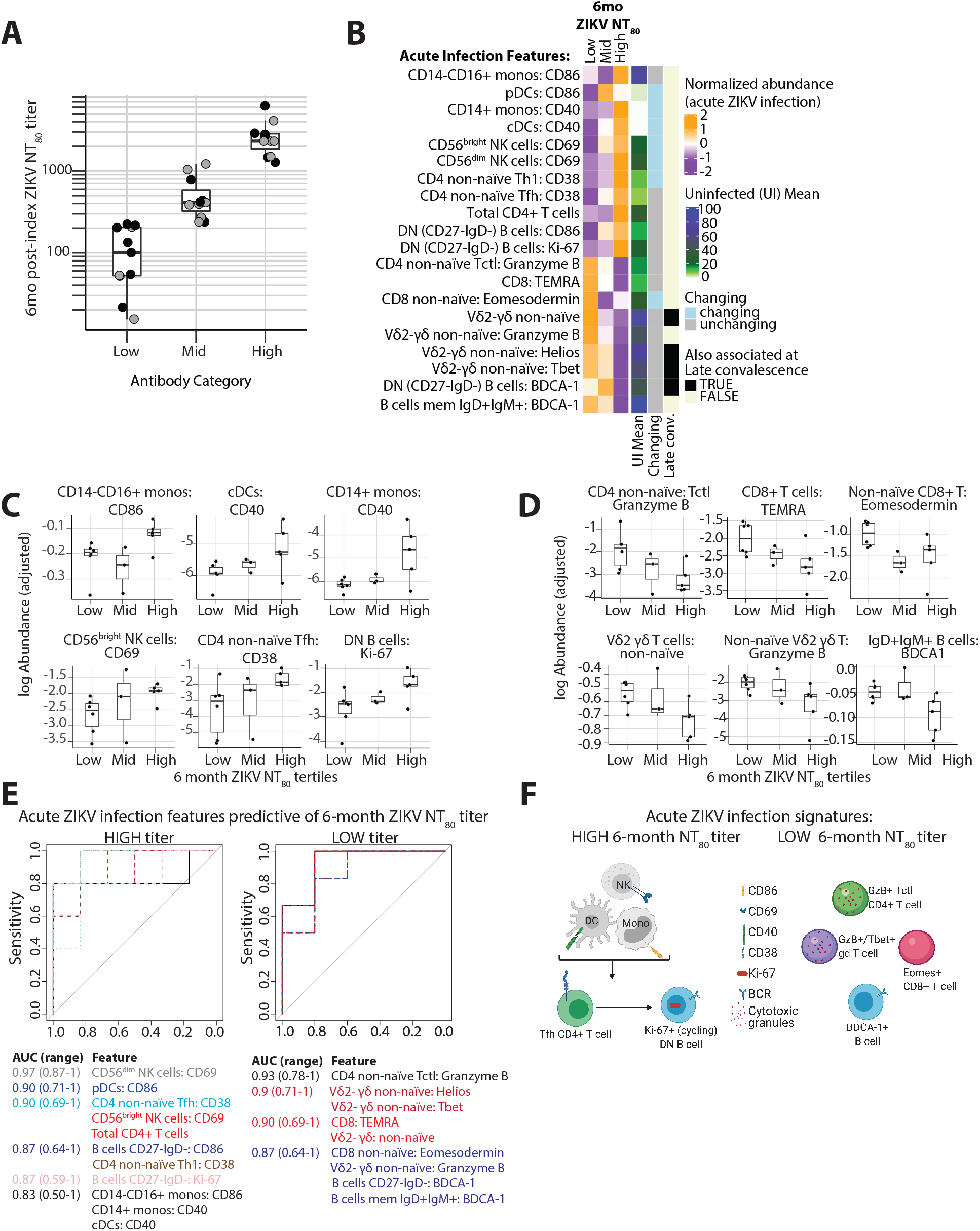
Distinct cellular immune signatures are associated with the development of high versus low ZIKV neutralizing antibody titers 6 months after infection. (A) Boxplot of ZIKV neutralizing antibody titers (NT80) measured approximately 6 months post-index visit in the overall REDS-III study participants (grey dots) and the sub-cohort studied here (black dots). Participants were divided into tertiles based on these values. (B) Heatmap showing z-score normalized abundance at the acute visit for cellular features that were significantly (p_adj<0.05) increased in high versus low 6-month NT80 titer participants at the acute timepoint. Also shown for each feature: Mean values in a cross-sectional uninfected (UI) control cohort; whether or not the abundance of the feature significantly changed across time between acute to convalescent infection; whether or not the abundance of the feature was also present at a significantly higher frequency (p_adj<0.05) in the same group (high-versus low-titer participants) at the late convalescent timepoint. Abundance (log-adjusted) of features during acute ZIKV infection associated with high (C) versus low (D) 6-month neutralizing antibody titers. (E) Receiver operating characteristic (ROC) curves for the cellular features from (B) associated with high (left) versus low (right) 6-month ZIKV neutralizing antibody titers. The area under the curve (AUC) value and 95% CI for the features corresponding to each curve are colored by AUC value. (F) Graphical illustration of cellular immune features from acute ZIKV infection associated with high versus low 6-month ZIKV neutralizing antibody titers.

To evaluate how our manually gated cellular features were coordinated during acute ZIKV infection, we initially focused on B cell and CD8+ T cell features that are known to be enriched for antigen-specific cells. We found that the frequencies of ASC B cells and CD38+HLA-DR+ non-naïve CD8+ T cells were positively correlated (Spearman’s r=0.61, p=0.01; **Fig. 5A**). To explore how innate and adaptive immune cellular features were more broadly co-regulated during acute ZIKV infection, we created a correlation heatmap of cellular features at the acute phase timepoint (including both cell type and phenotypic feature abundances; **Fig. S8**). This analysis revealed 279 positively and 66 negatively (p_adj<0.05) correlated pairs of features during acute ZIKV infection.

In order to characterize the pairs that were significantly correlated uniquely during acute ZIKV infection (and not in the uninfected state in the control participants), we determined whether the 95% confidence interval (CI) for each significant correlation during acute ZIKV infection did not contain (“unique” correlation, n=169) or did contain (“shared” correlation, n=176) the correlation value for the same feature pair in the cross-sectional uninfected samples (**Fig. 5B**). Compared to the “shared” correlations, the “unique” correlations during acute ZIKV infection had increased odds of being negatively correlated (odds of correlations being negative/positive amongst “unique” [49/120] versus “shared” [17/159] correlations; odds ratio [OR] 3.80 [95% CI: 2.03-7.42]). For example, uniquely in acute ZIKV infection, we observed negative correlations in the frequency of activated B cells versus CD4+ Tregs (r=-0.86, p_adj=0.002) and CD69+ CD56^dim^ NK cells versus Helios+ Vδ2-γδ T cells (r=-0.77, p_adj=0.03; **Fig. 5C, D**). Furthermore, compared to the “shared” correlations, the “unique” correlations were more likely to be correlations between features from different major landmark populations (odds of correlations being between different/same landmark populations amongst “unique” [105/64] versus “shared” [68/108] correlations; OR 2.60 [1.65-4.12]). The “unique” correlations were also more likely to be between (rather than within) adaptive and innate immune features (odds of correlations being between/within innate and adaptive features amongst “unique” [48/121] versus “shared” [22/154] correlations; OR 2.77 [1.54-5.10]). For example, we observed “unique” positive correlations between CD38+ pDCs and CD38+ Th1 CD4+ T cells (r=0.79, p_adj=0.03) and between the frequency of CD40+ cDCs and Ki-67+ double negative (DN; CD27-IgD-) B cells (r=0.78, p_adj=0.03; **Fig. 5E**).

Amongst phenotypic features, positive correlations present uniquely in acute ZIKV infection (total n=106) had increased odds of being between two different markers versus being between the same marker (odds of positive correlations being between different/same markers amongst “unique” [71/35] versus “shared” [66/74] correlations; OR 2.27 [1.31-3.98]). Of the list of significant positive correlations between different markers unique to acute ZIKV infection, the most common marker pair was CD40 and CD38 (n=5 edges) and the most frequently appearing markers included these and several other activation markers (CD38 [n=16 edges], CD40 [n=14 edges], ICOS [n=14 edges], and Ki-67 [n=11 edges]; **Fig. 5F**). In contrast to the “unique” pairs, the “shared” positive correlations present in both acute ZIKV infection and the uninfected state (which had increased odds of being between the same marker) featured subnetworks of the same phenotypic marker expressed on different cell types (e.g., CD69, CD38, CD71, and PD-1; depicted in **Fig. 5G**).

### Symptomatic ZIKV Infection is Associated with Greater Activation of Multiple Immune Cell Types

From the 17 participants enrolled prior to the development of IgM, we asked which manually gated populations at the acute infection timepoint differed were present at significantly different frequencies between participants who were symptomatic (≥3 symptoms at acute visit; n=8) or asymptomatic (n=9). Symptomatic participants during acute ZIKV infection had a higher frequency of multiple activated immune cell populations, with the strongest associations being amongst T cell (e.g., Ki-67+ γδ T cells, ICOS+ Tfh CD4+ T cells, HLA-DR+ non-naïve CD8+ T cells) and B cell features (e.g., IgD+IgM+ B cells expressing PD-1 and CD69). In contrast, asymptomatic individuals had a higher overall frequency of CD14+ and CD14-CD16+ monocytes, pDCs, Helios+ Tctl CD4+ T cells, and CD86+ transitional and DN B cells (**Fig. S9**).

### The Acute-phase Cellular Immune Response to ZIKV Infection Predicts Neutralizing Antibody Titers

We observed a large range in the titers of ZIKV neutralizing antibodies (NT_80_) that persisted several months after the resolution of acute infection (**Fig. 1B**). This prompted us to investigate which age- and sex-adjusted acute infection cellular immune features were associated with the development of a high versus low ZIKV NT_80_ titer 6 months after infection. Since prior exposure to DENV can be associated with significantly higher long-term ZIKV NT_80_ antibody titers ((*11*) and **Fig. S10A**), we excluded from this analysis 2 of the 17 early-stage participants (pre-IgM at the index visit, as described above) whose serologic profile at the index visit indicated that they had not previously been exposed to DENV. One early-infection stage participant who did not have samples collected past the acute phase visit was also excluded from this analysis (final n=14).

To our surprise, we did not observe a correlation between the frequencies of ASCs and HLA-DR+CD38+ CD8+ T cells at the acute timepoint and the level of ZIKV NT_80_ at 6 months post-infection (**Fig. S10B, C**). Therefore, we decided to more broadly explore associations between the 6-month ZIKV NT_80_ titers and all of the cellular immune features that we had characterized at the acute timepoint. To do this, we first subdivided the larger DENV-exposed REDS-III cohort (total n=33; **Fig. 6A**) into three groups according to their 6-month ZIKV NT_80_ titers (measured a median of 181 days after index visit [range 160-196 days]). Of note, there was no significant difference in the age or sex distribution between the three tertiles as determined by one-way ANOVA (p=0.31) and a chi-square test of independence (p=0.53), respectively. We used the tertile boundaries from the full REDS-III cohort to assign each of the participants from our cohort to the low (n=6, <230), mid-range (n=3, 230-1240) or high (n=5, >1240) ZIKV NT_80_ titer group. We then used an exact permutation test to determine which cellular features at the acute timepoint had significantly different frequencies between individuals in the low versus the high 6-month ZIKV NT_80_ tertiles.

We found that high levels of ZIKV neutralizing antibody titers 6 months post-infection were associated with a significantly higher frequency of a total of 11 cellular features during acute infection, including multiple activated cell types: (1) CD14-CD16+ monocytes and pDCs that express the costimulatory molecule CD86, (2) CD14+ monocytes and cDCs that express the costimulatory molecule CD40, (3) NK cells that express CD69, (4) Th1 and Tfh CD4+ T cells that express CD38 (as well as a higher overall frequency of total CD4+ T cells), and (5) DN B cells that express CD86 and Ki-67 (**Fig. 6B**, with select populations shown in **6C**). When measured in the uninfected state, the group of features associated with high antibody titers tend to be present at lower frequencies compared to the group of features associated with low antibody titers (see lighter green colors in “Uninfected [UI] Mean” column). The high antibody titer features also trended towards having more features that significantly decreased in frequency between acute infection and convalescent timepoints (light blue in “Changing” column; 6 changing of 11 total features in the low titer group versus 1/9 in the high titer group; OR=8.53, p=0.07). Together, these data suggest that high 6-month ZIKV NT_80_ titers are associated with the dynamic expansion during the acute phase of infection of activated cell populations that are present at lower frequencies in the absence of infection.

In contrast, individuals with low titers of neutralizing ZIKV antibodies 6 months after infection had an immune signature during acute infection defined by higher frequencies of T cell populations with a cytotoxic profile (higher Granzyme B expression in Tctl CD4+ T cells, a larger TEMRA population in CD8+ T cells and higher Eomesodermin expression non-naïve CD8+ T cells, a higher overall frequency of non-naïve Vδ2-γδ T cells and a higher frequency of non-naïve Vδ2-γδ T cells that express Granzyme B, T-bet and Helios) as well as a higher frequency of DN B cells and IgD+IgM+ memory B cells that express BDCA1 (**Fig. 6B**, with select populations shown in **6D**). Unlike the acute infection cellular features associated with high 6-month NT_80_ titers, most of the acute infection cellular features associated with low 6-month NT_80_ titers were larger cell populations in the uninfected state (see darker green/blue colors in the “Uninfected Mean” column) and most (n=8 of 9) were not dynamically regulated over the course of ZIKV infection (grey in “Changing” column).

As a proxy for understanding how features of an individual’s pre-infection immune state may relate to the development of high or low durable ZIKV NT_80_ titers, we again used an exact permutation test to ask which cellular features measured after the resolution of infection at the late convalescent timepoint were associated with 6-month ZIKV NT_80_ titers. Interestingly, we noted overlap with a subset of features associated with low, but not high, 6-month ZIKV NT_80_ titers (black in “Late Convalescence” column). Collectively, these data suggest that low 6-month ZIKV NT_80_ titers may be associated with an “immune set-point” characterized by a higher frequency of cytotoxic-differentiated T cell populations that are not dynamically regulated during acute ZIKV infection. Because a higher frequency of cytotoxic-differentiated T cells can be associated with a history of infection with other viruses, in particular cytomegalovirus (CMV), and a positive CMV serostatus can be associated with impaired response to vaccination (*53–55*), we asked whether CMV seropositivity was associated with the development of low ZIKV NT_80_ titers. We did not observe this association (p=0.29).

To determine whether the acute ZIKV infection cellular immune features associated with the development of high versus low 6-month ZIKV NT_80_ titers could potentially be used to predict these outcomes, we performed a receiver operating characteristic (ROC) analysis. This analysis revealed that all of the acute infection features associated with high or low antibody titers also reliably predicted these two outcomes in this cohort (minimum area under the curve [AUC]=0.833; **Fig. 6E**). Several of the features in the low antibody titer signature with the highest predictive power (e.g., non-naïve Vδ2-γδ T cells that express Tbet, Helios, AUC=0.900) did not significantly change over the course of ZIKV infection and were also increased in the low-titer individuals during the late convalescent timepoint, suggesting they may represent an immune “set-point” that predisposes to the generation of lower neutralizing antibody titers in response to ZIKV infection. In fact, phenotypic features of Vδ2-γδ T cells exhibited the most unique negative correlations of any cell type (**Fig. S10D**), suggesting their connectivity with other immunological parameters that may dictate such a “set-point”. Furthermore, during acute ZIKV infection, the cellular feature that had the highest predictive power of a high 6-month neutralizing antibody titer (CD56^dim^ NK cells:CD69, AUC=0.967) was significantly negatively correlated with the frequency of the low NT_80_ titer-associated Helios+ Vδ2-γδ T cell population **(Fig. 5D**). This observation suggests that the cellular immune features predictive of high versus low 6-month ZIKV neutralizing antibody titers may truly represent distinct immune states (**Fig. 6F**).

## DISCUSSION

We present here the first deep characterization of cellular immune features that are dynamically regulated during acute ZIKV infection in human adults. We leveraged mass cytometry as a systems immunology approach to characterize activated cell populations across the innate and adaptive immune system during an acute infection in humans. Our goal was to understand how these features are coordinated during the acute phase of infection and to identify immune signatures associated with the development of durable high titers of ZIKV neutralizing antibody at 6 months post-infection. From our longitudinal cohort of individuals found to be viremic at the time of blood donation, we found that acute ZIKV infection is characterized by the dynamic and coordinated expression of several activation markers across different cell types in the innate and adaptive immune system. High levels of ZIKV neutralizing titers 6 months after infection were associated with transiently higher frequencies of multiple activated innate immune subsets, as well as activated Tfh and proliferating DN B cells during acute infection. On the other hand, low levels of 6-month ZIKV neutralizing titers were associated with immune features that had characteristics suggestive of a cytotoxic-skewed immune “set-point.”

Our study adds to a small but growing body of literature that has characterized diverse cellular immune phenotypes in the acute phase of infection with other viral pathogens in humans, including dengue (*46, 56, 57*), Ebola (*49*), influenza (*47, 58*), chikungunya (*59*), HIV (*60–63*), and SARS-CoV-2 (*31, 48, 64–66*) viruses. In prior studies of ZIKV infection, acute infection has been associated with increased production of several plasma cytokines (*34, 35*), activation of some innate immune cell types (*38*) and, in children, an increase in the frequency of monocyte populations that are also a target for viral infection *in vivo* (*37, 45*). Here, we applied mass cytometry panels that we specifically designed to identify cell types and phenotypic markers that we reasoned would likely be dynamically regulated during acute viral infection. Because acute ZIKV infection can be associated with a reduction in absolute numbers of lymphocytes in the peripheral blood (*38*), we were surprised that acute ZIKV infection did not alter the frequency of most major landmark cellular populations. On the other hand, we did observe significant changes in the frequency of 128 of 286 manually gated features and 38 of 90 unsupervised cell clusters over the course of infection, many of which represented small populations of cells that expressed multiple activation markers and were transiently increased at the acute timepoint. Furthermore, we observed by PCA that the cellular immune signature of most participants followed a similar trajectory across time from the acute to the convalescent phase. PCA revealed that sampling time during acute infection was significantly associated with the distance and starting point of each participant’s trajectory in PCA space, a proxy for the longitudinal dynamic immune signatures of the samples. These analyses revealed the power of our targeted and detailed CyTOF panels to identify dynamically regulated features on small populations of cells during acute ZIKV infection. Our findings also suggested to us that, in this acute infection cohort, it was critical to account for sampling time (*66*).

Leveraging the power of our longitudinal cohort to overcome the statistical limitations of high inter-individual variability in human immune phenotypes, we identified several notable populations of immune cells that are relatively increased during acute ZIKV infection. Amongst innate immune cells, we observed an expansion of intermediate CD14+CD16+ monocytes during acute infection as has also been noted in a pediatric ZIKV infection cohort (*37, 45*), and we found that this population was highly activated in the acute phase. Additionally, we identified a cluster of monocytes that were present at a higher frequency during acute infection (c49) in which the co-expression of markers that denote proliferation versus activation versus differentiation state appear to be distinctly co-regulated. Amongst adaptive immune cells, we observed an expansion of HLA-DR+CD38+ non-naïve CD8+ T cells, which has been observed in other acute viral infections (*31, 46, 47*), and we noted that this cell population contained four distinct clusters of cells that co-express different combinations of activation markers. Acute ZIKV infection was also associated with activation of Th1 and Tctl T cell CD4+ T cell subsets. Finally, using gating strategies to identify populations of B cells enriched for antigen-specific cells in other infections (*49, 67*), we identified an expansion of Tbet+ ASCs during acute ZIKV infection. Our study is the first to describe the diverse and coordinated activation of cellular immune responses during acute ZIKV infection in human adults. Similar patterns of activation across innate and adaptive immune cell types have also been observed in SARS-CoV-2 infection using high-dimensional flow and mass cytometry as well as single cell RNA sequencing (*31, 64, 68*), suggesting substantial conservation in the immune response to distinct acute viral infections.

One strength of our study is the inclusion of PBMC samples from participants in a cross-sectional ZIKV-uninfected control cohort collected based at the same blood donation sites prior to the onset of the ZIKV epidemic. This cohort allowed us to identify how activation of different cell populations is uniquely coordinated during acute ZIKV infection. Compared to the baseline correlations that exist in the uninfected state, we found that during acute ZIKV infection there was an increased odds that the correlations were between features derived from different landmark populations and across the innate and adaptive immune system. This included correlations that may reflect the immune populations that need to be coordinated to mount a productive immune response in this infection (e.g., positive correlations in the proportion of CD38+ pDCs and CD38+ Th1 CD4+ T cells, or CD40+ cDCs and Ki-67+ DN B cells). Analyzing different immune parameters, a coordinated immune response during acute SARS-CoV-2 infection has been associated a with mild compared to severe disease (*69*). Taken together with our data, these studies suggest that mass cytometry and other systems immunology approaches that can evaluate the degree to which early immune responses across different cell types are coordinated may be useful to identify markers and pathways that predict disease outcome.

Animal models of ZIKV infection suggest that both cellular and humoral immunity may be important for protection from re-infection (*12, 13, 70*). The titer of ZIKV neutralizing antibodies is likely a critical component of protective immunity in humans and is a key target for ZIKV vaccines in clinical development (*71*). Furthermore, low titers of ZIKV neutralizing antibodies may theoretically put people at risk for enhancing disease upon infection with the cross-reactive flavivirus, DENV (*72*). Six months following infection, participants across our cohort had a greater than 100-fold difference in the titers of their neutralizing antibodies. Other than a positive association with prior DENV serostatus observed here and in other studies (*11*), little is known about what immune parameters may predispose some individuals to maintain higher versus lower ZIKV neutralizing antibody titers.

Using our window into immune cell dynamics during the acute phase of infection, we observed that high 6-month ZIKV neutralizing antibody titers were associated with a higher frequency of several activated innate and adaptive immune cell features that mostly were present at low frequencies in uninfected state and many of which significantly declined in frequency between acute and convalescent infection. Furthermore, ROC analysis indicated that each of the cell features associated with high- or low-neutralizing antibody titers also reliably predicted the assignments to these groups. Interestingly, as has been in observed in SARS-CoV-2 infection (*31*), the frequency of ASCs during acute infection did not correlate with antibody levels in convalescence. However, many of the cellular features that we found were associated with high neutralizing antibody titers have plausible roles in augmenting a productive B cell response. For example, CD86 expression on pDCs and monocytes and CD40 expression on cDCs and monocytes can mediate enhanced antigen presentation to and priming of helper CD4+ T cells, IFNγ produced by activated Th1 cells or NK cells can promote B cell activation, and activated Tfh CD4+ T cells can provide direct help to differentiating B cells. A higher frequency of these cellular immune features during the acute phase of natural ZIKV infection thus may reflect the activity of an immune response that supports the formation of durably high levels of neutralizing antibodies. Further investigation will be useful for elucidating mechanisms that underlie the variable immunogenicity profiles of different vaccine platforms for ZIKV and potentially other viral infections, such as SARS-CoV-2.

In contrast to the cellular immune features associated with high ZIKV neutralizing antibody titers, we observed a very different acute infection cellular immune signature amongst individuals who had low ZIKV neutralizing antibody titers 6 months after infection. Low 6-month ZIKV neutralizing antibody titers were associated with a higher frequency of T cells with cytotoxic differentiation features, several of which did not dynamically change over the course of infection and were also at higher frequency in these individuals at the late convalescent timepoint, after cell frequencies had returned to levels indistinguishable from the uninfected state. This pattern of expression suggested to us that a cytotoxic immune “set-point” may define the low titer group. In general, a more cytotoxic-skewed T cell compartment can be a sign of immune senescence, which can in turn be associated with a reduced capacity to generate functional antigen-specific responses after vaccination (*53, 55*). The fact that we observed a negative correlation between two features that reliably predicted either the high- or the low-neutralizing antibody titer outcome (CD69+ CD56^dim^ NK cells and Helios+ Vδ2- γδT cells, respectively) may indicate that these two immune signatures truly represent distinct immune states.

Our study has some important limitations. First, while we had the advantage of working with clinical samples collected in the context of a cohort with detailed clinical annotation, this was a small study constrained by the number of participants in the larger REDS-III cohort who had appropriate PBMC and serum samples available. Although we have made an effort to control for the variance introduced by sampling time, it is not possible to align participants according to the date they were infected in natural infection cohorts. Our study included only otherwise healthy individuals who presented for volunteer blood donation, and thus is not representative of the broader population and, in particular, does not include pregnant individuals or infants, who are key populations affected by this viral infection and whose immune responses are likely to be quite different from our study population. Our mass cytometry panels were designed to characterize cellular features that we reasoned would be impacted by ZIKV infection. However, the changes we observed are limited by the markers we chose to study. Nonetheless, we feel this approach has some important advantages to other systems immunology approaches. For example, with our detailed phenotypic panels, we had the resolution to precisely identify small populations of cells that change in frequency over time, a phenomenon that may be missed or poorly characterized with bulk or even single-cell RNA-sequencing or with CyTOF panels built to monitor broad immune cell populations. Finally, we sought to correlate cellular immune features from acute infection with the formation of ZIKV neutralizing antibody titers. While it is likely that neutralizing antibodies play a key role in immunologic protection from ZIKV and other acute viral infections (*73*), a titer that correlates with protection in humans has not been identified (*16*) and other antibody functions (*74*) and/or other types of immune responses may be critical to providing robust protection.

Our study is the first to provide an in-depth characterization of the cellular immune response to acute ZIKV infection in human adults and to relate this response to the development of high- or low-titers of durable neutralizing antibodies. We have identified cellular features that suggest immune responses that may contribute to the generation of protective immune responses in this natural viral infection in humans. Our approach offers a powerful tool to test whether these features predict immunogenicity of different vaccine platforms for ZIKV and potentially other viral infections, such as SARS-CoV-2, for which protection is thought to be largely due to the activity of neutralizing antibodies.

## MATERIALS AND METHODS

### Study Design

This study was designed as a retrospective cohort study of a sub-group of participants in the observational prospective REDS-III cohort of individuals who were found to have PCR-documented acute ZIKV infection at the time of blood donation. Participants were included if they had PBMC sampling available at acute infection. Participants were excluded if the interval between index visit (blood donation) and the first PBMC blood draw was greater than two weeks (n=1 participant excluded based on this criteria). Mass cytometry experiments were performed over the course of five separate experiments, with normalization between experiments performed as outlined below.

### Cohort and Participant Characteristics

We characterized the cellular immune response to acute and resolving ZIKV infection in the peripheral blood of 25 otherwise healthy adults (negative for HIV, hepatitis B and C infections) who had viremic ZIKV infection between 2015-2016 at the time of blood donation in Puerto Rico (“index” visit). Participants were selected from the prospective REDS-III cohort (see **Table S2** for demographic and clinical characteristics) (*42*). Peripheral blood mononuclear cells (PBMCs) from these ZIKV+ individuals were sampled longitudinally at up to three intervals after the index visit: (1) within 5-12 days after the index visit (“acute”; median 8 days), (2) within 15 to 27 days after index visit (“early convalescence”; median 21 days), and (3) within 85-189 days after index visit (“late convalescence”; median 91 days; see **Fig. 1A** for sampling schema). ZIKV viral load and antibody measurements were performed at these and additional timepoints (see **Fig. 1B**). Participants were asked about the presence of 6 symptoms at each visit (fever, rash, joint or bone pain, body or muscle pain, painful or red eyes, headache). For the purposes of this study, participants were designated as “symptomatic” if they reported three or more symptoms present at the acute visit (first PBMC sampling timepoint). Two groups of ZIKV-uninfected (“ZIKV-”) individuals were also included in our analyses. The ZIKV-participants were recruited from a larger cohort of blood donors based at the same blood donation clinics as the ZIKV+ participants (with the same sample processing protocols), with samples collected in the years prior to the onset of the ZIKV epidemic in Puerto Rico. Samples from the first ZIKV-cohort (single timepoint, n=8) were processed and run on the mass cytometer in parallel with the samples from the ZIKV+ individuals. Samples from the longitudinal ZIKV-cohort (n=6 participants across three timepoints) were run separately. All samples were obtained with appropriate informed consent and ethics committee approval of the University of California San Francisco.

### ZIKV Viral Load, ZIKV and DENV Antibody Levels, and ZIKV Neutralizing Antibody Levels

ZIKV viral load, antibody levels, and ZIKV and DENV neutralizing antibody measurements were performed as described previously (*9, 41*). In brief, ZIKV viral load was measured by quantitative PCR. Anti-Zika virus IgM and IgG were measured by antibody-capture ELISA using recombinant ZIKV antigen kindly provided by the US Centers for Disease Control and Prevention (CDC) and as previously described (*75, 76*). ZIKV neutralizing titers were measured using a ZIKV reporter viral particle neutralization titration assay (Integral Molecular, Philadelphia, PA) (*77*), and index donations were tested for pre-existing DENV IgG with the Detect IgG ELISA (InBios; Seattle, WA).

### PBMC Preparation and Staining for Mass Cytometry

Whole peripheral blood was collected at the clinical sites, shipped overnight at ambient temperature to Vitalant, San Francisco, CA, USA, where they were processed and cryopreserved within 24 h of collection and then stored in liquid nitrogen as previously described (*42*). PBMCs were thawed, and only samples with >70% viability were used for analysis (most were >90% viable after thawing by the Muse Cell Analyzer [Millipore Sigma, Burlington, MA, USA]) (*78, 79*). We stained 2-4 million cells per panel in two mass cytometry panels (see **Table S2** for antibody clones and metals), following a previously published protocol (*44*) with the following modifications. Briefly, we marked dead cells by incubating the samples for one minute with 25mM Cisplatin (Sigma-Aldrich, St. Louis, MO, USA) in phosphate buffered saline (PBS) plus EDTA, performed surface staining with metal-tagged antibodies in PBS with 0.5% bovine serum albumin (BSA) for 30 minutes at room temperature, fixed and permeabilized cells following manufacturer’s instructions for the eBioscience Foxp3/Transcription Factor Staining Buffer Set (Thermo Fisher Scientific, Waltham, MA, USA), barcoded samples using mass-tag cellular barcoding reagents diluted in Maxpar Barcode Perm Buffer (Fluidigm, South San Francisco, CA, USA) as described previously (*44*), combined up to twenty barcoded samples into a single tube, performed intracellular staining with antibodies diluted in eBioscience Foxp3/Transcription Factor kit perm wash (Thermo Fisher Scientific), fixed cells in freshly prepared 2% paraformaldehyde (Electron Microscopy Sciences, Hatfield, PA, USA) in the presence of a DNA intercalator (*80*), and then washed and ran cells on the Fluidigm CyTOF 2 mass Cytometer within one week of staining.

### Mass Cytometry Data Processing

#### Data Quality Control

Following data acquisition, the FCS files were normalized across experiments using bead standards and the data normalization algorithm using the R package ‘premessa’ (https://github.com/ParkerICI/premessa). The live cell events were debarcoded using a single-cell debarcoding algorithm (*81*) and we analyzed >25,000 (mostly >50,000) cells per sample. From the individual sample files, normalization beads were excluded based on Ce140 and Eu153 signal, single cell events were identified based on Ir191 DNA signal measured against event length, and CD45- or Pt195+ dead cells were excluded. Potential batch effects were minimized by including samples from the same individual in the same experiment. Gating was performed using CellEngine (CellCarta, Montreal, Canada).

#### Manual Gating

Traditional hierarchical gating was applied to identify 12 “landmark” immune populations: CD14+ “classical” monocytes, CD14-CD16+ “non-classical” monocytes, classical and plasmacytoid dendritic cells [cDC and pDC, respectively], basophils, CD56^bright^ and CD56^dim^ natural killer cells, regulatory CD4+ T cells, non-regulatory CD4+ T cells, CD8+ T cells, γδ T cells as stained by either a pan-γδ T cell receptor (TCR) antibody or an antibody that only recognizes γδ T cells with the Vδ2 chain (see **Fig. S1** for gating strategy) as well as well-defined adaptive immune subsets (see **Fig. S2** for the identity of these populations). Within each of the “parent” cell types, we manually gated positive and negative populations of biologically relevant phenotypic markers from the two mass cytometry panels (see **Table S3** for markers assessed on each “parent” population). For each of the parent cell types, we only included phenotypic markers for which we could clearly gate a positive population above background antibody staining levels.

#### Clustering by Statistical SCAFFoLD

We generated SCAFFoLD maps using the statisticalScaffold R package (available at https://github.com/SpitzerLab/statisticalScaffold). As described previously (*43, 44*), using all of the live CD45+ leukocytes collected across participants and timepoints for each staining panel, we applied an unsupervised clustering algorithm based on the CLARA clustering algorithm to partition cells into a user-defined number of clusters (100 clusters per staining panel). We excluded Ki-67 and Granzyme B to avoid having functional markers cluster cells across cell types together. Landmark populations were gated as outlined in **Fig. S1** (for cluster analysis, NK cells were treated as one population). We next generated force-directed graphs (SCAFFoLD maps) to visualize the association of each cluster with its likely parent landmark population. We excluded from our downstream analysis clusters that contained <20 cells in >80% of samples (12 clusters in Panel 1, 2 clusters in Panel 2) as well as clusters that contained cells that did not have the expected expression of classical landmark population (e.g., we excluded a cluster of cells that clustered with the CD8+ T cells but appeared to co-express the B cell marker, CD19 and may potentially represent doublets [median 0.09% of total CD8+ T cells at the acute timepoint]; all together, these 9 clusters in Panel 1 and 7 clusters in Panel 2 comprised 0.08% and 0.13% of the total live population at the acute timepoint). Cell clusters were thus determined to be “reliably” assigned to landmark cell populations if they were not excluded based on these criteria and if they were identified in Panel 1 for innate immune cells (total 17 classical and 3 non-classical monocyte, 9 NK cell, 4 cDC, 1 pDC clusters) and B cells (total 14 clusters) and Panel 2 for T cell phenotypes (total 6 CD4+ Treg, 20 non-Treg CD4+, 14 CD8+, and 2 non-Vδ2 γδ T cell clusters). In the SCAFFoLD maps depicted, a representative map from one participant at timepoint 1 is shown.

### Statistical Analysis

#### Change in manually gated population and cell cluster frequencies over time

To measure the change in abundance of manually gated cell features (e.g., landmark and sub-landmark populations and populations expressing individual phenotypic markers) and cell clusters, the frequency of each feature (expressed as a % of the parent population) was log transformed with a constant factor of 1/10e6 or 1/10e3, respectively. Log-transformed values were adjusted for participant age and sex using a linear regression and the residuals (log-adjusted abundance) were used in downstream analyses. The change over time for the log-adjusted feature abundance between the “acute,” “early convalescent” and “late convalescent” visits was assessed using linear mixed effect (LME) model fit with log-transformed days since index visit as a fixed effect and participant ID as a random effect. The p values for each group of features were adjusted for multiple testing correction by Benjamini Hochberg with an FDR cutoff of 5% for a significant effect of time since index visit on feature abundance. For 95% confidence interval graphs, line graphs were generated in R using the packages grid (base), ggplot (3.2.1), gridExtra (2.3), and lattice (0.20-38). The 95% confidence intervals for the median values were calculated by bootstrapping with 1000 iterations.

#### PCA Analysis

The log-adjusted manually gated features that were present across all ZIKV infected and cross-sectional uninfected samples (281 of 324 total features in the dataset) were used for principal component analysis with the function PCA (parameters: “scale.unit = TRUE”, ncp = 5) from the R package FactoMineR (1.42). The samples were visualized in PCA space with PC1 and PC2 values as the coordinates using ggplot2 (3.2.1) in R.

#### Heatmaps

Heatmaps were made in R using the package ComplexHeatmap (2.1.1). For the manually gated features and cluster features summary heatmaps, the row/column orders, respectively, were determined using the R package seriation (1.2.8) with the travelling salesperson problem (TSP) method.

#### Network correlation analysis

Pairwise Spearman correlations were calculated on the on the log-adjusted feature abundances from samples at the acute visit for participants who were previously exposed to Dengue and in an early stage of infection (pre-IgM at the time of Index visit). The p values were adjusted with the Benjamini-Hochberg method. The 95% confidence intervals for the correlation in the infected samples for each pairwise feature comparison was calculated using bootstrapping with 1000 iterations. For each significant correlation (p adjusted<0.05), the correlation was categorized as shared with the uninfected cohort if the correlation value in the uninfected cohort fell within the 95% confidence interval from the infected samples or had the same sign as the infected correlation and a magnitude greater than the 95% confidence interval magnitude maximum. Fisher’s exact test was used to determine odds ratio for correlations being unique to ZIKV as the exposure (versus being shared with the uninfected) and the indicated correlation attribute as the outcome.

#### Antibody and Symptom Associations

The NT_80_ titers at the 6-month timepoint of the DENV-exposed, ZIKV+ individuals from the larger REDS III cohort were classified into antibody tertiles. For the symptom associations, we used acute visit samples from participants who had not yet formed IgM at the index visit (n=17) and for the antibody associations, we further subset participants who were DENV-exposed who had 6-month NT_80_ titers available (n=14). Exact permutation tests were used to test for significant differences in the log-adjusted cellular features between samples from participants in the high versus low tertiles (n=5 in high group and n=6 in low group) and between samples from participants classified as symptomatic or asymptomatic (n=8 for symptomatic and n=9 for asymptomatic). The association between CMV IgG seropositivity and 6-month NT_80_ titers was assessed using the Wilcoxon Rank Sum test based on a larger subset of REDS-III study participants for whom CMV serostatus were available (n=10 CMV+, n=23 CMV-).

#### Antibody Associations Predictive Modelling

We used pROC (v 1.17.0.1) to plot ROC curves with log-adjusted feature abundance at the acute visit as the predictor and 6-month NT_80_ antibody titer category (e.g., “High” or “Low”) as the response for each participant. The 95% CI for the AUC values were computed with the default “DeLong” method.

## Supplementary Materials

**Fig. S1.** CyTOF gating strategy.

**Fig. S2.** Principal Component Analysis (PCA) of uninfected participants.

**Fig. S3.** Landmark and sub-landmark population abundance in acute and convalescent ZIKV infection.

**Fig. S4.** Landmark population frequencies by manual gating versus clustering.

**Fig. S5.** cDC, pDC and NK cell features impacted by acute ZIKV infection.

**Fig. S6.** Treg and γδ T cell features impacted by acute ZIKV infection.

**Fig S7.** B cell dynamics in ZIKV infection.

**Fig S8.** Immune cell features correlated during acute ZIKV infection.

**Fig S9.** Association of acute ZIKV infection cellular immune features with the development of symptoms.

**Fig. S10.** Characteristics associated with ZIKV neutralizing antibody titers 6 months after infection.

**Table S1.** Mass cytometry antibodies used.

**Table S2.** Study participant clinical characteristics.

**Table S3.** Summary of phenotypic markers assessed on each cell type for manual gating analysis.

**Data file S1.** Raw data table of study participant ZIKV viral load, total IgM and IgG levels, and neutralizing antibody titers.

**Data file S2.** Raw data tables of manually gated immune cell features (as a % of CD45+ live cells for landmark features, and as a % of parent populations for all others) and the frequency of SCAFFoLD clusters (as a % of parent population) for all study participants and timepoints.

**Data file S3:** Statistical testing results: linear mixed effect modeling, cellular features that are significantly correlated in acute ZIKV infection and/or in the uninfected state, antibody and symptom association permutation testing results.

## Acknowledgements

We thank Stanley Tamaki for expert assistance with mass cytometry and the Parnassus Flow Core Facility. Figures 1A and 6F created with BioRender.com.

## Funding

National Institutes of Health grant DP5OD023056 (MHS)

National Institutes of Health grant T32AI060530 (RLR) National Institutes of Health grant K23AI134327 (RLR)

National Institutes of Health grant T32GM007618 (EEM)

National Institutes of Health grant F30CA257291 (EEM)

National Institutes of Health grant P30DK063720 (Parnassus Flow Core)

National Institutes of Health grant S10OD018040 (Parnassus Flow Core)

National Institutes of Health grant S10OD021822 (Parnassus Flow Core)

Department of Health and Human Services Contract HHSN26820110001I (MPB)

MHS is a Chan Zuckerberg Biohub Investigator.

## Author contributions

Conceptualization: RLR, PMO, MHS

Cohort management and participant selection: MPB, MS

Methodology (CyTOF staining): RLR, PMO, EL, CPS, IT

Methodology (serology, neutralization titers): GS, PJN

Data Analysis: EEM, RLR, MHS, PMO

Data Visualization: EEM

Manuscript writing: RLR, EEM, MHS (with input from all authors)

Funding acquisition: MHS

Supervision: RLR, MHS, PWH, MEF

## Data and materials availability

Raw data files will be made publicly available upon manuscript acceptance. All data generated from manual gating and clustering analysis are provided in supplemental tables.

