## Supplementary Figures for "Distinct cellular immune signatures in acute Zika virus infection are associated with high or low persisting neutralizing antibody titers"

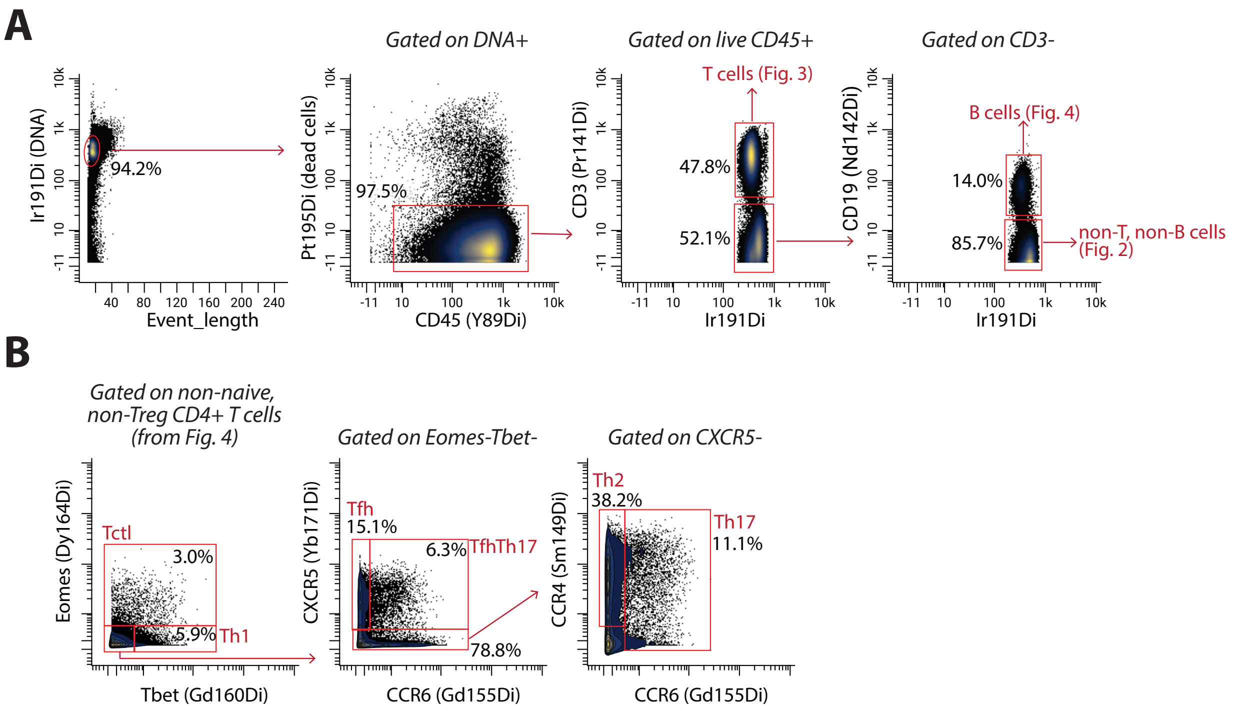


**Figure S1. CyTOF gating strategy**
Immune cell (A) landmark population and (B) non-naïve CD4+ T cell sub-population gating strategy.

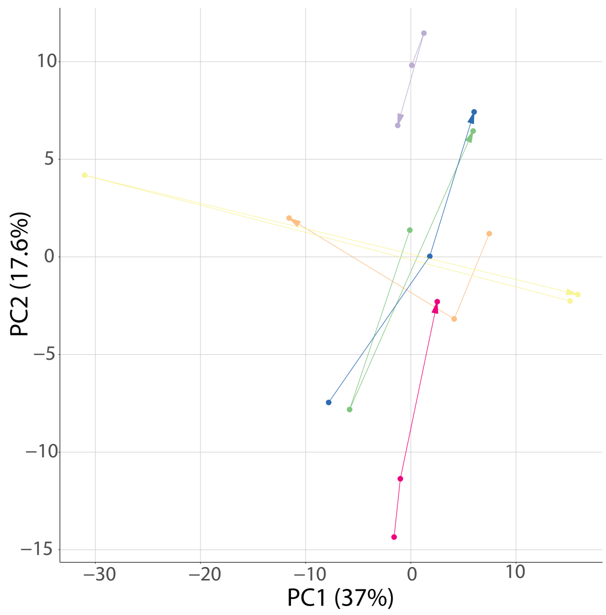


**Figure S2: Principal Component Analysis (PCA) of uninfected participants.**
PCA representation of all manually gated parameters measured on PBMCs from ZIKV-uninfected control participants at longitudinal timepoints.

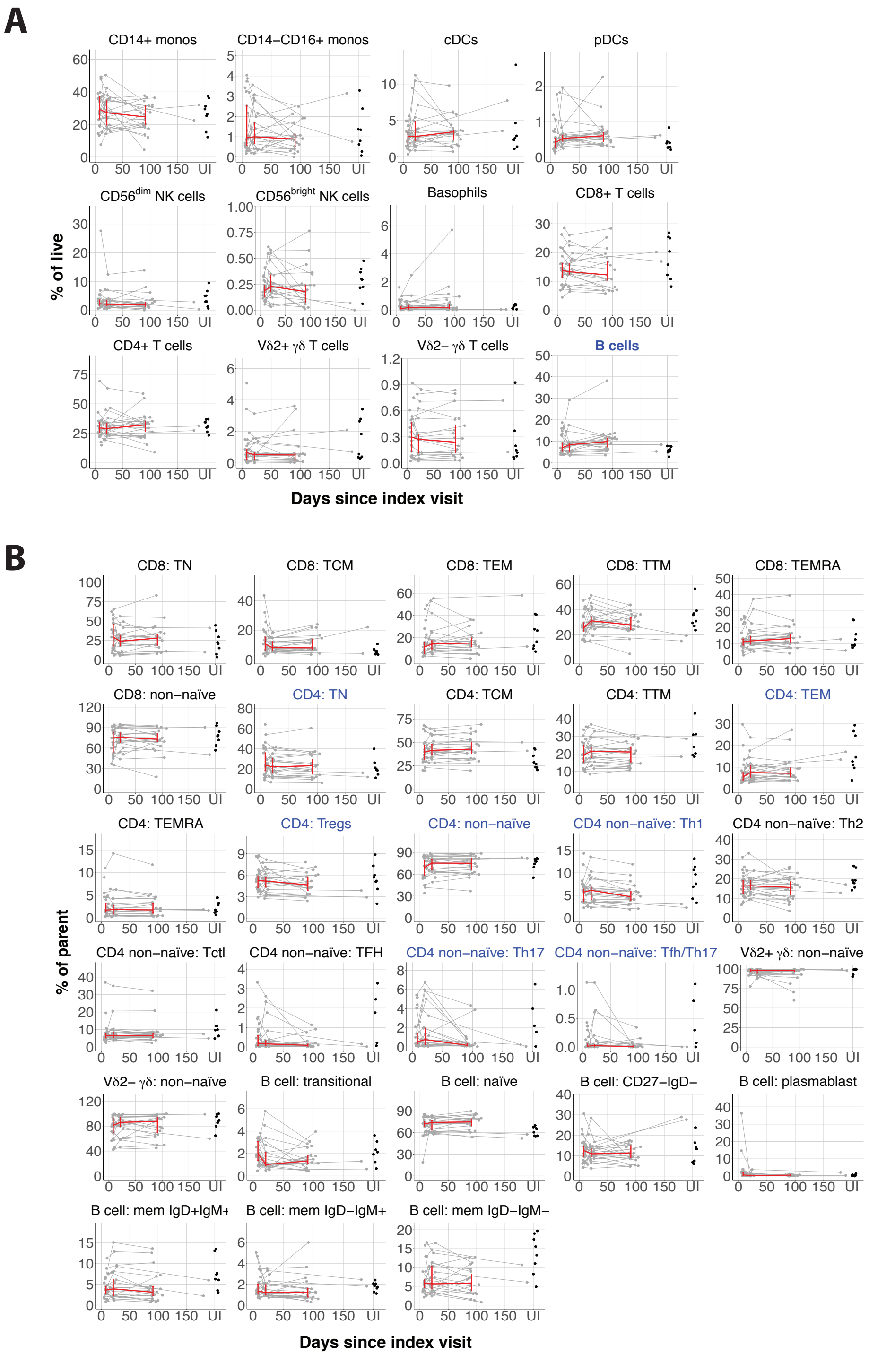


**Figure S3. Landmark and sub-landmark population abundance in acute and convalescent ZIKV infection.** Line plots of frequency of indicated (A) landmark cell type and adaptive immune subset for each participant versus time since index visit. Red line connects median values at each sampling timepoint with error bars for +/- 95% CI. Scatterplot for feature abundance from cross-sectional uninfected (UI) cohort shown on the far right. Features with p_adj< 0.05 have blue colored titles. p_adj values obtained by LME model fit with Benjamini-Hochberg FDR correction.

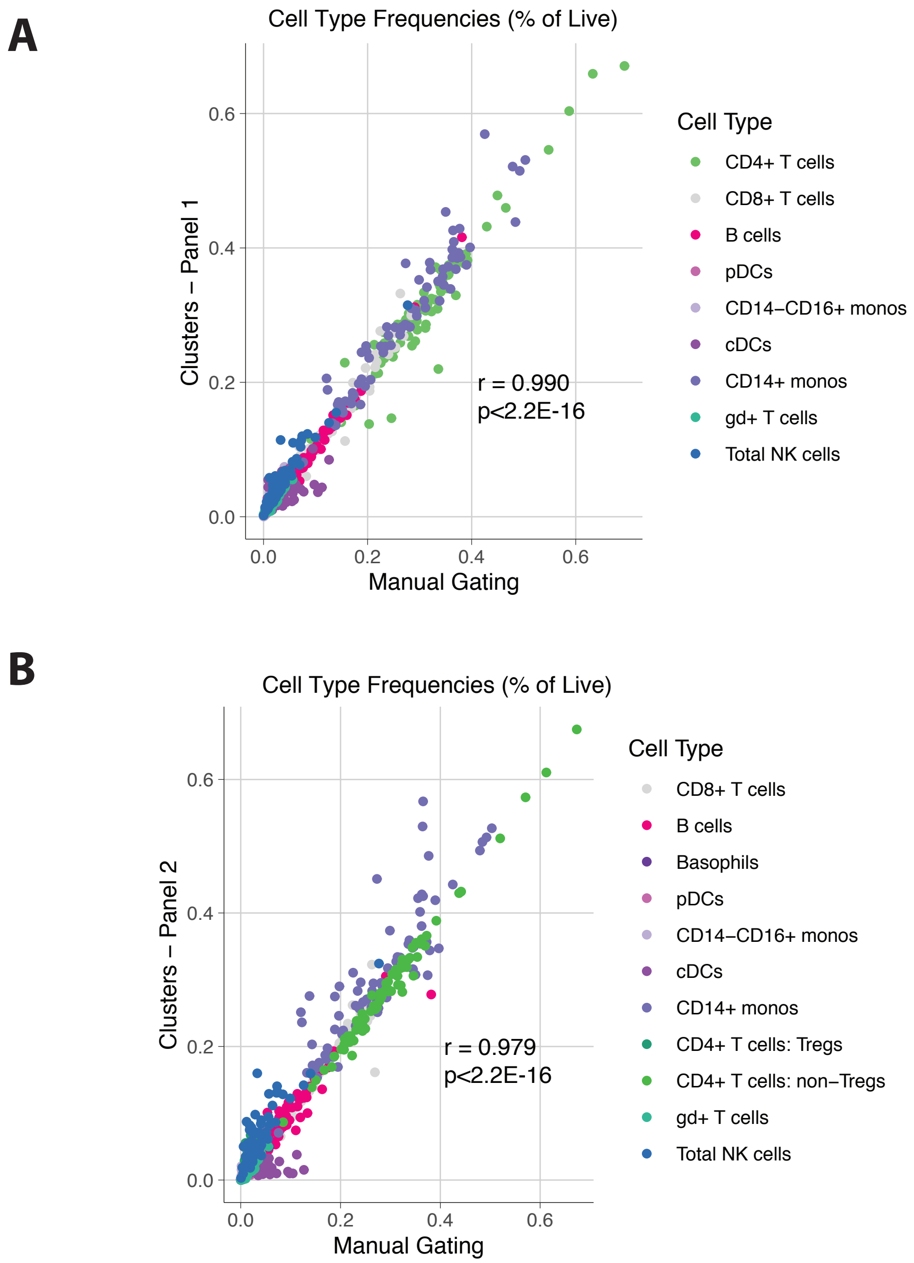


**Figure S4. Landmark cell population frequencies by manual gating vs clustering.**
High concordance in landmark cell population frequencies as measured by manual gating versus SCAFFoLD clustering analysis.

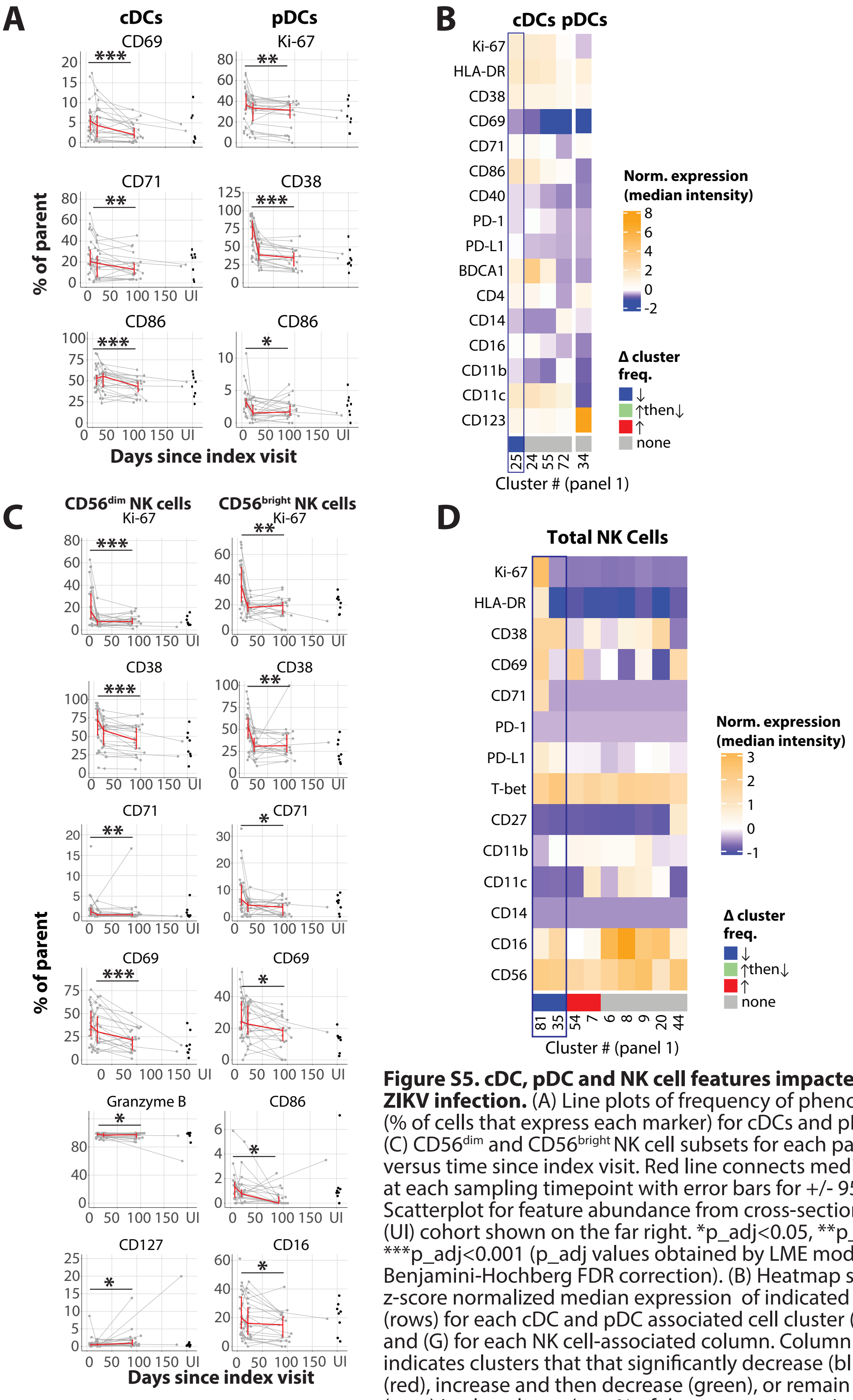


**Figure S5. cDC, pDC and NK cell features impacted by acute ZIKV infection.** (A) Line plots of frequency of phenotypic features (% of cells that express each marker) for cDCs and pDCs and (C) CD56$^{dim}$ and CD56$^{bright}$ NK cell subsets for each participant versus time since index visit. Red line connects median values at each sampling timepoint with error bars for +/- 95% CI. Scatterplot for feature abundance from cross-sectional uninfected (UI) cohort shown on the far right. *p_adj<0.05, **p_adj<0.01, ***p_adj<0.001 (p_adj values obtained by LME model fit with Benjamini-Hochberg FDR correction). (B) Heatmap showing z-score normalized median expression of indicated markers (rows) for each cDC and pDC associated cell cluster (column) and (G) for each NK cell-associated column. Column annotation indicates clusters that that significantly decrease (blue), increase (red), increase and then decrease (green), or remain unchanged (grey) in abundance (as a % of the parent population; p_adj<0.05).

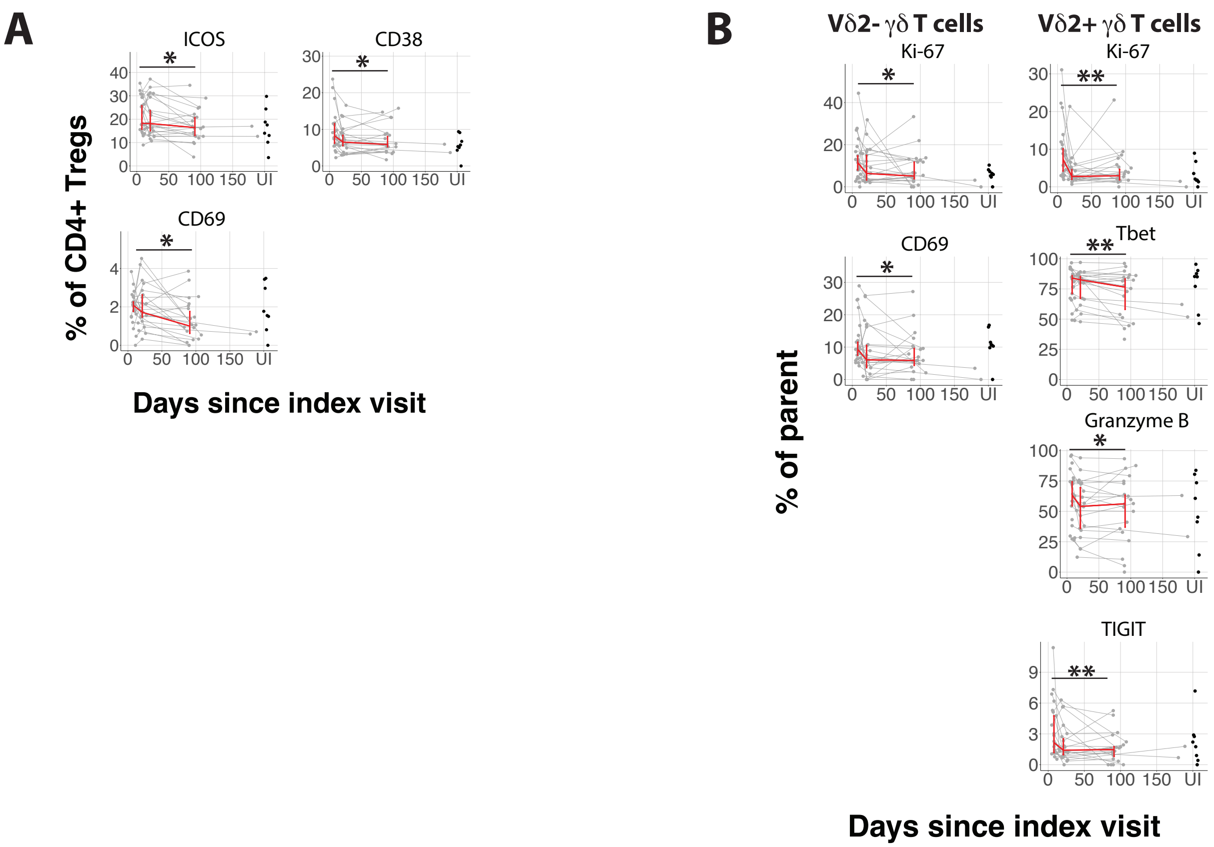


**Figure S6. Treg and gd T cell features impacted by acute ZIKV infection.**
(A) Line plots of frequency of phenotypic features (% of cells that express each marker) for Tregs and (B) gd T cells subsets for each participant versus time since index visit. Red line connects median values at each sampling timepoint with error bars for +/- 95% CI. Scatterplot for feature abundance from cross-sectional uninfected (UI) cohort shown on the far right. *p_adj<0.05, **p_adj<0.01, ***p_adj<0.001 (p_adj values obtained by LME model fit with Benjamini-Hochberg FDR correction).

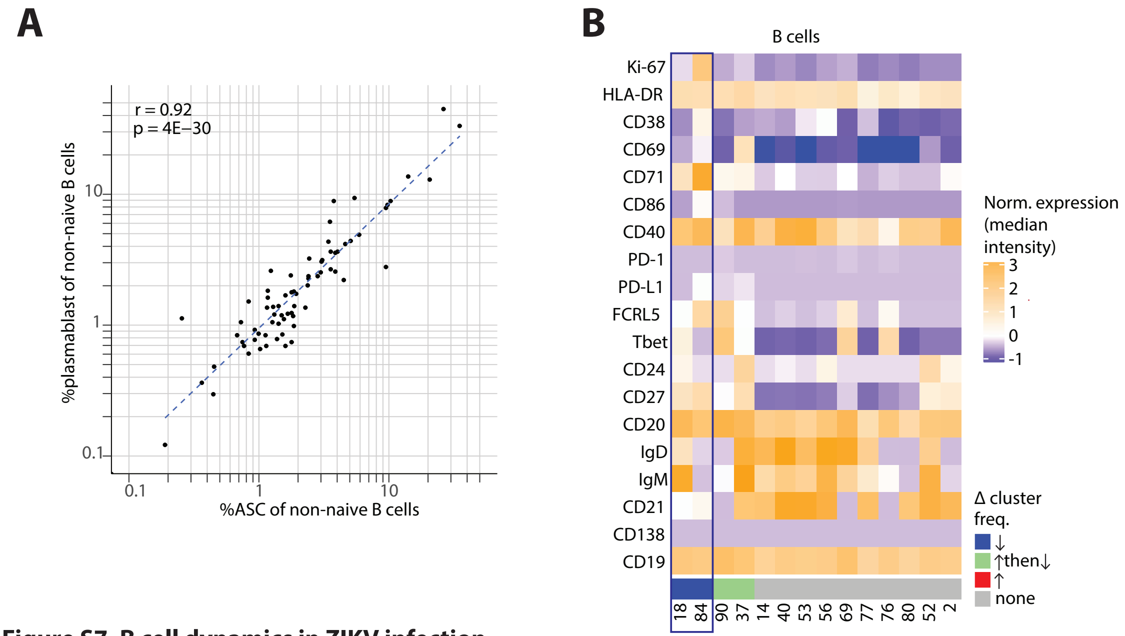


**Figure S7. B cell dynamics in ZIKV infection.**
(A) Correlation plot between the frequency of plasmablast and ASC B cell populations (as a % of non-naïve B cells; Spearman's r with regression line). (B) Phenotype (z-scored median expression of each marker) of B cell clusters that significantly decrease (blue), increase (red), increase and then decrease (green), or remain unchanged (grey) in abundance (as a % of the total B cell population; p_adj<0.05).

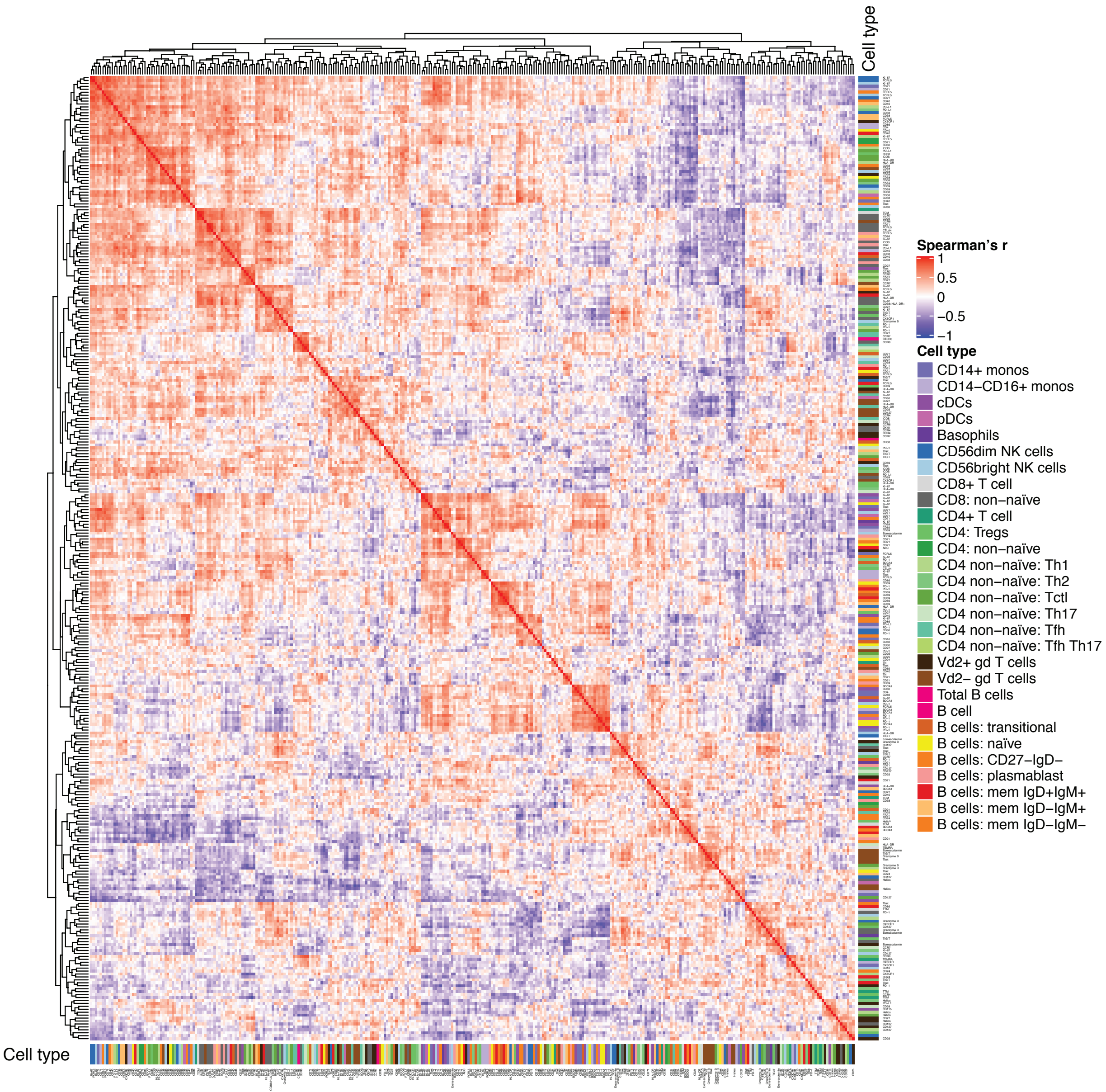


**Figure S8. Immune cell features correlated during acute ZIKV infection.**
Correlation heatmap depicting Spearman's correlation values (no significance cut-off) of all manually gated features from acute ZIKV infection (n=17 pre-IgM participants).

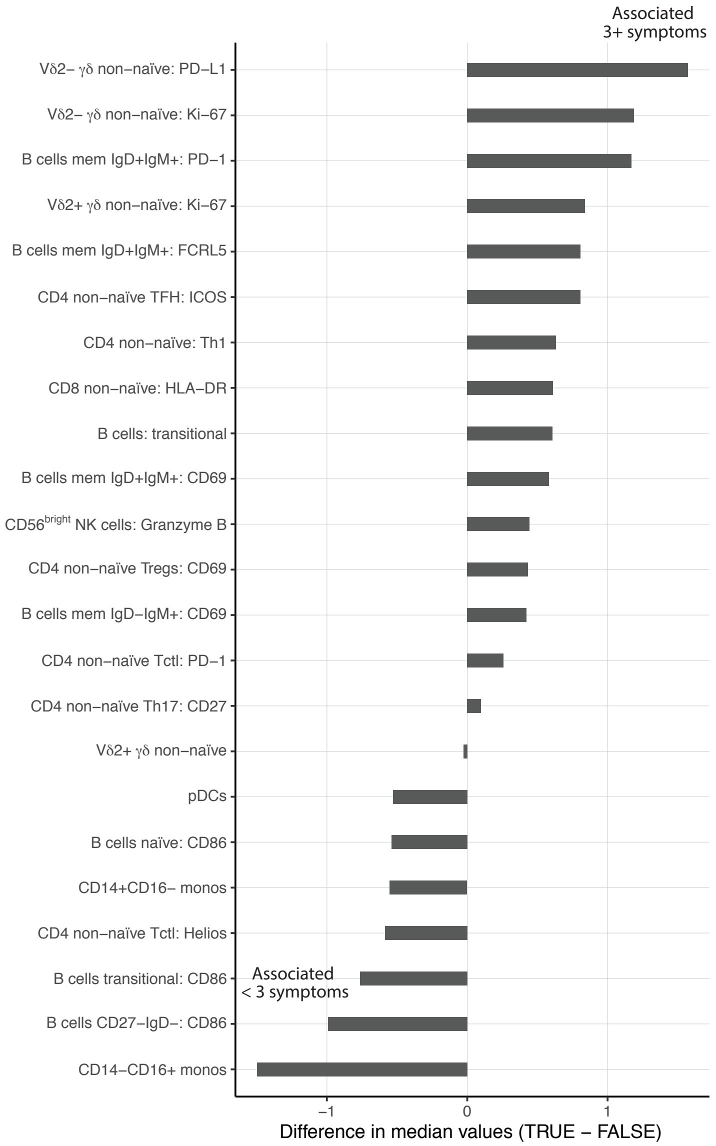


**Figure S9. Association of acute ZIKV infection cellular immune features with the development of symptoms.** Barplot of difference in median values at the acute visit for cellular immune features whose abundance during acute ZIKV infection is associated with the development of 3 or more symptoms reported at the acute visit (=TRUE; $p<0.05$ by exact permutation test).

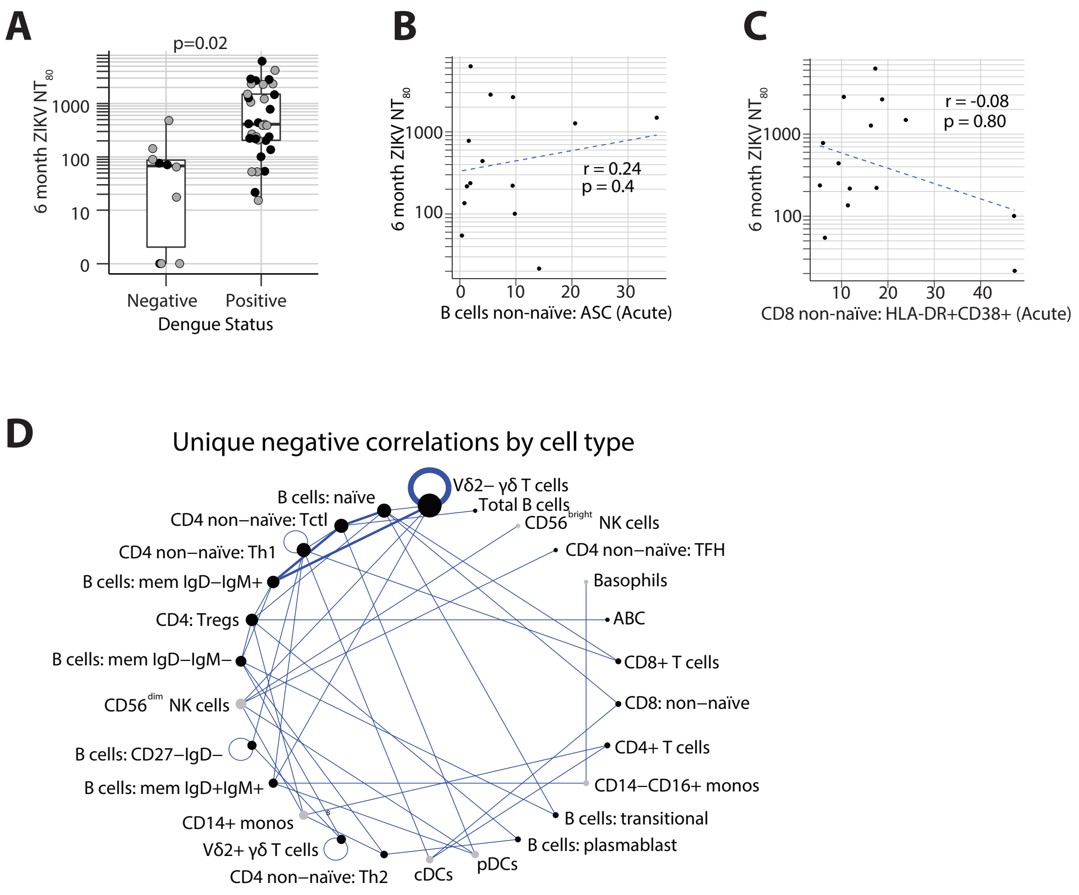


**Figure S10. Characteristics associated with ZIKV neutralizing antibody titers 6 months after infection.** (A) Difference in 6-month ZIKV $NT_{80}$ between individuals with or without evidence of prior DENV infection at index visit (Wilcoxon Rank Sum test). Individuals from our sub-cohort are colored black and individuals from the larger REDSIII cohort are colored grey. Scatterplots showing 6-month ZIKV $NT_{80}$ titers versus the frequency of (B) ASC B cells or (C) non-naïve CD8+ T cells co-expressing HLA-DR and CD38 at the acute timpoint (Spearman's correlation). (D) Circular network graph of negative correlations unique to acute ZIKV infection. Size of node indicates number of correlations and edge thickness is proportional to the number of correlations between features classified by the indicated nodes. Nodes indicate cell type. Correlations between features of the same cell type are indicated with a circular line segment and nodes from adaptive immune cell types are colored black and from innate immune cell types are colored grey.

**PANEL 1**

| Antigen | Metal | Clone | Vendor | Catalogue # | In-house conjugated Abs: [Ab] (mg/mL) | In-house conjugated Abs: [Staining] (ug/mL); Fluidigm: vol/100uL |
|---|---|---|---|---|---|---|
| CD45 | 89 | HI30 | Fluidigm | 3089003B | | 0.5 |
| CD14 | 113 | M5E2 | Biolegend | 301802 | 0.500 | 3.000 |
| CD123 | 115 | 6H6 | Biolegend | 306002 | 0.5 | 0.750 |
| CD33 | 139 | WM53 | Biolegend | 303402 | 0.5 | 0.750 |
| CD38 | 140 | HIT2 | Biolegend | 303502 | 0.5 | 3.000 |
| CD3 | 141 | UCHT1 | Biolegend | 300402 | 0.5 | 1.000 |
| CD19 | 142 | H1B19 | Biolegend | 302202 | 0.5 | 1.500 |
| CXCR3 | 143 | G025H7 | Biolegend | 353702 | 0.5 | 1.500 |
| CD11b | 144 | ICRF44 | Biolegend | 301302 | 0.5 | 1.5 |
| CD4 | 145 | RPA-T4 | Biolegend | 300502 | 0.5 | 0.250 |
| CD8 | 146 | RPA-T8 | Biolegend | 301002 | 0.5 | 0.500 |
| CD11c | 147 | Bu15 | Biolegend | 337202 | 0.5 | 0.375 |
| CD16 | 148 | 3G8 | Biolegend | 302001 | 0.3 | 3.000 |
| CD138 | 149 | DL-101 | Biolegend | 352302 | 0.5 | 1.500 |
| CD21 | 151 | Bu32 | Biolegend | 313502 | 0.5 | 1.500 |
| gdTCR | 152 | 11F2 | Fluidigm | 3152008B | | 0.5 |
| CD45RA | 153 | HI100 | Biolegend | 304102 | 0.5 | 0.750 |
| CD40 | 154 | 5C3 | Biolegend | 334302 | 0.5 | 6.000 |
| PDL1 | 156 | 29E.2A3 | Biolegend | 329702 | 0.5 | 1.500 |
| CD69 | 157 | FN50 | Biolegend | 310902 | 0.5 | 0.750 |
| CD27 | 158 | O323 | Biolegend | 302802 | 0.5 | 1.500 |
| Tbet | 160 | 4B10 | Biolegend | 644802 | 0.5 | 1.500 |
| CTLA4 | 161 | 14D3 | Fluidigm | 3161004B | | 0.25 |
| CD80 | 162 | 2D10.4 | Fluidigm | 3162010B | | 0.5 |
| CD86 | 163 | IT2.2 | Biolegend | 305401 | 0.5 | 0.750 |
| CD24 | 165 | MI5 | Biolegend | 311102 | 0.5 | 3.000 |
| NKG2D | 166 | ON72 | Fluidigm | 3166016B | | 1 |
| FCRL5 | 167 | 509f6 | Biolegend | 340302 | 0.5 | 3.000 |
| Ki67 | 168 | B56 | Fluidigm | 3168007B | | 1 |
| CD71 | 169 | CY1G4 | Biolegend | 334102 | 0.3 | 0.750 |
| IgD | 170 | CY1G4 | Biolegend | 334102 | 0.3 | 0.750 |
| CD20 | 171 | 2H7 | Biolegend | 302302 | 0.3 | 1.500 |
| BDCA1 | 172 | L161 | Biolegend | 331502 | 0.3 | 0.375 |
| IgM | 173 | MHM-88 | Biolegend | 314502 | 0.5 | 1.500 |
| HLA-DR | 174 | L243 | Biolegend | 307602 | 0.5 | 1.500 |
| PD-1 | 175 | EH12.2H7 | Biolegend | 329902 | 0.5 | 1.500 |
| CD56 | 176 | HCD56 | Fluidigm | 3176008B | | 0.5 |

**PANEL 2**

| Antigen | Metal | Clone | Vendor | Catalogue # | In-house conjugated Abs: [Ab] (mg/mL) | In-house conjugated Abs: [Staining] (ug/mL); Fluidigm: vol/100uL |
|---|---|---|---|---|---|---|
| CD45 | 89 | HI30 | Fluidigm | 3089003B | 0.5 | 0.5 |
| CD14 | 113 | M5E2 | Biolegend | 301802 | 0.500 | 3.000 |
| CD123 | 115 | 6H6 | Biolegend | 306002 | 0.5 | 0.750 |
| CD33 | 139 | WM53 | Biolegend | 303402 | 0.5 | 0.750 |
| CD38 | 140 | HIT2 | Biolegend | 303502 | 0.5 | 3.000 |
| CD3 | 141 | UCHT1 | Biolegend | 300402 | 0.5 | 1.000 |
| CD19 | 142 | H1B19 | Biolegend | 302202 | 0.5 | 1.500 |
| CXCR3 | 143 | G025H7 | Biolegend | 353702 | 0.5 | 1.500 |
| CD11b | 144 | ICRF44 | Biolegend | 301302 | 0.5 | 1.5 |
| CD4 | 145 | RPA-T4 | Biolegend | 300502 | 0.5 | 0.250 |
| CD8 | 146 | RPA-T8 | Biolegend | 301002 | 0.5 | 0.500 |
| CD11c | 147 | Bu15 | Biolegend | 337202 | 0.5 | 0.375 |
| CD16 | 148 | 3G8 | Biolegend | 302001 | 0.3 | 3.000 |
| CCR4 | 149 | 205410 | R+D | MAB1567 | 0.5 | 3.000 |
| OX40 | 150 | A019D5 | Biolegend | 351302 | 0.5 | 3.000 |
| ICOS | 151 | C398.4A | Biolegend | 313539 | 0.5 | 1.500 |
| gdTCR | 152 | 11F2 | Fluidigm | 3152008B | | 0.5 |
| CD45RA | 153 | HI100 | Biolegend | 304102 | 0.5 | 0.750 |
| CX3CR1 | 154 | 2A9-1 | Biolegend | 341602 | 0.2 | 0.375 |
| CCR6 | 155 | G034E3 | Biolegend | 353402 | 0.3 | 1.500 |
| PDL1 | 156 | 29E.2A3 | Biolegend | 329702 | 0.5 | 1.500 |
| CD69 | 157 | FN50 | Biolegend | 310902 | 0.5 | 0.750 |
| CD27 | 158 | O323 | Biolegend | 302802 | 0.5 | 1.500 |
| Vd2 | 159 | B6 | Biolegend | 331402 | 0.2 | 0.750 |
| Tbet | 160 | 4B10 | Biolegend | 644802 | 0.5 | 1.500 |
| CTLA4 | 161 | 14D3 | Fluidigm | 3161004B | | 0.25 |
| FOXP3 | 162 | PCH101 | Fluidigm | 3162011a | | 0.25 |
| EOMES | 164 | WD1928 | ThermoFisher | 14-4877-82 | 0.5 | 3.000 |
| CD127 | 165 | A019D5 | Biolegend | 351302 | 0.5 | 1.500 |
| TIGIT | 166 | A15153G | Biolegend | 372702 | 0.5 | 1.500 |
| CCR7 | 167 | G043H7 | Biolegend | 353202 | 0.5 | 1.500 |
| Ki67 | 168 | B56 | Fluidigm | 3168007B | | 1 |
| CD25 | 169 | 2A3 | Fluidigm | 3169003B | | 1 |
| CXCR5 | 171 | RF8B2 | Fluidigm | 3171014B | | 0.125 |
| Helios | 172 | 22F6 | Biolegend | 137202 | 0.5 | 1.500 |
| Granzyme B | 173 | GB11 | BioRad | MCA2120 | 0.5 | 1.500 |
| HLA-DR | 174 | L243 | Biolegend | 307602 | 0.5 | 1.500 |
| PD-1 | 175 | EH12.2H7 | Biolegend | 329902 | 0.5 | 1.500 |
| CD56 | 176 | HCD56 | Fluidigm | 3176008B | | 0.5 |

**Table S1.** Mass cytometry antibodies used.

**ZIKV-infected**

| Participant # | Age | Sex | Pre-IgM at index visit | DENV exposed | >=3 symptoms at acute visit | Maximum ZIKV NT80 titer | 6mo ZIKV NT80 titer |
|---|---|---|---|---|---|---|---|
| 1 | 52 | Male | Y | Y | TRUE | 355.1 | 54.3 |
| 2 | 24 | Male | N | N | FALSE | 84.4 | 0 |
| 3 | 22 | Male | N | Y | TRUE | 9719.1 | 2894.4 |
| 4 | 62 | Male | Y | Y | FALSE | 1069.7 | 217.1 |
| 5 | 43 | Male | Y | N | FALSE | NA | NA |
| 6 | 54 | Male | N | Y | FALSE | 13952.6 | 419.9 |
| 7 | 51 | Male | Y | Y | FALSE | 2782.7 | 221.7 |
| 8 | 46 | Male | Y | Y | TRUE | 688 | 21.6 |
| 9 | 49 | Male | N | N | FALSE | 1474.1 | 71.8 |
| 10 | 37 | Male | Y | Y | FALSE | 37872.3 | 6285.7 |
| 11 | 46 | Male | Y | Y | TRUE | 3086.1 | 100.2 |
| 12 | 36 | Female | N | Y | TRUE | 4543 | 202.9 |
| 13 | 43 | Male | Y | Y | TRUE | 3405.1 | 1483.7 |
| 14 | 42 | Male | Y | Y | TRUE | 2238.7 | 134.8 |
| 15 | 43 | Female | N | N | Unknown | 1153 | 77.1 |
| 16 | 28 | Female | N | N | Unknown | NA | NA |
| 17 | 27 | Female | Y | N | TRUE | NA | NA |
| 18 | 53 | Male | N | N | FALSE | 1198.7 | NA |
| 19 | 25 | Female | Y | Y | FALSE | NA | NA |
| 20 | 71 | Male | Y | Y | FALSE | 2091.4 | 440.9 |
| 21 | 67 | Male | Y | Y | FALSE | 3752.8 | 237.6 |
| 22 | 24 | Male | Y | Y | FALSE | 2206.1 | 2660.2 |
| 23 | 56 | Female | Y | Y | TRUE | 9412.2 | 1278.7 |
| 24 | 44 | Male | Y | Y | FALSE | 23236.4 | 2828.5 |
| 25 | 21 | Female | Y | Y | TRUE | 1490.7 | 779.1 |

**ZIKV-uninfected**

| Participant # | Age | Sex |
|---|---|---|
| 26 | 49 | Male |
| 27 | 32 | Female |
| 28 | 51 | Female |
| 29 | 60 | Male |
| 30 | 53 | Male |
| 31 | 58 | Male |
| 32 | 42 | Female |
| 33 | 22 | Male |
| 34 | 32 | Female |
| 35 | 20 | Male |
| 36 | 40 | Male |
| 37 | 54 | Male |
| 38 | 44 | Male |
| 39 | 20 | Male |

**Table S2.** Study participant clinical characteristics.

| Cell type | Ki-67 | HLA-DR | CD38 | CD69 | CD71 | CD86 | CD16 | CD40 | ICOS | CTLA-4 | TIGIT | PD-1 | PD-L1 | Granzyme B | Tbet | Eomesodermin | Helios |
|---|---|---|---|---|---|---|---|---|---|---|---|---|---|---|---|---|---|
| CD14+ monocytes | x | | | x | x | x | x | x | | | | x | x | | x | | |
| CD14-CD16+ monocytes | x | | | x | x | x | | x | | | | x | | | x | | |
| cDCs | x | x | | x | x | x | | x | | | | x | | | x | | |
| pDCs | x | | x | | x | x | | | | | | x | | | | | |
| CD56dim NK cells | x | x | x | x | x | x | | | | | x | x | | x | x | x | |
| CD56bright NK cells | x | x | x | x | x | x | x | | | | x | x | | x | x | x | |
| Basophils | x | | | | x | | | | | | | | | | | | |
| CD8: non-naïve | x | x | x | x | x | | | x | x | x | x | x | x | x | x | x | x |
| CD4: Tregs | x | x | x | x | | | | | x | x | x | x | | | | | x |
| CD4: non-naïve | | | | | x | | | | | | | | | | | | |
| CD4 non-naïve: Th1 | x | x | x | | | | | | x | | x | x | x | x | | | x |
| CD4 non-naïve: Th2 | x | x | x | | | | | | x | | x | x | x | | | | x |
| CD4 non-naïve: Tfh | x | x | x | | | | | | x | | | x | | | | | |
| CD4 non-naïve: Th17 | x | x | x | | | | | | | | x | x | | | | | |
| CD4 non-naïve: Tctl | x | x | x | | | | | | x | | x | x | x | x | | | x |
| Vd2+ gd T cells | x | x | x | x | | | | | | | x | x | x | x | x | x | x |
| Vd2- gd T cells | x | x | x | x | | | | | | | x | x | x | x | x | x | x |
| Total B cells | | | | | | | | | | | | | | | | | |
| B cells: transitional | x | | x | x | x | x | | x | | | | x | | | x | | |
| B cells: naïve | x | | x | x | x | x | | x | | | | x | | | x | | |
| B cells: CD27-IgD- | x | | x | x | x | x | | x | | | | x | | | x | | |
| B cells: plasmablast | x | | x | x | x | x | | x | | | | x | | | x | | |
| B cells: mem IgD+IgM+ | x | | x | x | x | x | | x | | | | x | | | x | | |
| B cells: mem IgD-IgM+ | x | | x | x | x | x | | x | | | | x | | | x | | |
| B cells: mem IgD-IgM- | x | | x | x | x | x | | x | | | | x | | | x | | |

| Cell type | CD25 | FCRL5 | CD21 | CD127 | CCR4 | CCR6 | CXCR5 | CX3CR1 | BDCA1 | CD4 | CD11b | CD27 | CCR7 | CD24 | total |
|---|---|---|---|---|---|---|---|---|---|---|---|---|---|---|---|
| CD14+ monocytes | | x | | | | | | x | x | x | | | | | 13 |
| CD14-CD16+ monocytes | | x | | | | | | x | x | x | | | | | 11 |
| cDCs | | | | | | | | | x | | x | | | | 10 |
| pDCs | | | | | | | | | | | | | | | 5 |
| CD56dim NK cells | | x | | x | | | | | | | | x | | | 14 |
| CD56bright NK cells | | x | | x | | | | | | | | x | | | 15 |
| Basophils | | | | | | | | | | | | | | | 2 |
| CD8: non-naïve | x | x | | x | x | x | | x | | | | x | x | | 23 |
| CD4: Tregs | | | | x | x | x | | | | | | x | x | | 14 |
| CD4: non-naïve | | x | | | | | | | | | | | | | 2 |
| CD4 non-naïve: Th1 | x | | | x | | | | x | | | | x | x | | 14 |
| CD4 non-naïve: Th2 | x | | | x | | | | | | | | x | x | | 12 |
| CD4 non-naïve: Tfh | x | | | x | | | | | | | | x | x | | 9 |
| CD4 non-naïve: Th17 | x | | | x | | | | | | | | x | x | | 9 |
| CD4 non-naïve: Tctl | x | | | x | | | | x | | | | x | x | | 14 |
| Vd2+ gd T cells | x | | | x | x | x | | x | | | | x | x | | 18 |
| Vd2- gd T cells | x | | | x | x | x | | x | | | | x | x | | 18 |
| Total B cells | | | | | | | x | | | | | | | | 1 |
| B cells: transitional | | x | x | | | | | | x | | | | | | 11 |
| B cells: naïve | | x | x | | | | | | x | | | | | x | 12 |
| B cells: CD27-IgD- | | x | x | | | | | | x | | | | | x | 12 |
| B cells: plasmablast | | x | x | | | | | | x | | | | | | 11 |
| B cells: mem IgD+IgM+ | | x | x | | | | | | x | | | | | x | 12 |
| B cells: mem IgD-IgM+ | | x | x | | | | | | x | | | | | x | 12 |
| B cells: mem IgD-IgM- | | x | x | | | | | | x | | | | | x | 12 |
| | | | | | | | | | | | | | | SUM: | 286 |

**Table S3.** Summary of phenotypic markers assessed on each cell type for manual gating analysis.
